# Variational phylodynamic inference using pandemic-scale data

**DOI:** 10.1101/2022.02.10.479891

**Authors:** Caleb Ki, Jonathan Terhorst

## Abstract

The ongoing global pandemic has sharply increased the amount of data available to researchers in epidemiology and public health. Unfortunately, few existing analysis tools are capable of exploiting all of the information contained in a pandemic-scale data set, resulting in missed opportunities for improved surveillance and contact tracing. In this paper, we develop the variational Bayesian skyline (VBSKY), a method for fitting Bayesian phylodynamic models to very large pathogen genetic data sets. By combining recent advances in phylodynamic modeling, scalable Bayesian inference and differentiable programming, along with a few tailored heuristics, VBSKY is capable of analyzing thousands of genomes in a few minutes, providing accurate estimates of epidemiologically relevant quantities such as the effective reproduction number and overall sampling effort through time. We illustrate the utility of our method by performing a rapid analysis of a large number of SARS-CoV-2 genomes, and demonstrate that the resulting estimates closely track those derived from alternative sources of public health data.

## 1 Introduction

The COVID-19 pandemic has demonstrated an important supporting role for phylogenetics in epidemiology and public health, while also creating unforeseen technical and methodological challenges. As the first global public health event to occur in an era of ubiquitous gene sequencing technology, the pandemic has resulted in a data explosion of unprecedented proportions. GISAID, a worldwide repository of SARS-CoV-2 genomic data, currently has over 7.5M samples, with contributions from almost every country (Elbe and Buckland-Merrett, 2017; van Dorp et al., 2021). A phylogenetic representation of this database is believed to be the largest ever constructed (Turakhia et al., 2021a). Existing phylogenetic methods, which were developed and tested on datasets orders of magnitude smaller, are inadequate for pandemic-scale analysis, resulting in missed opportunities to improve our surveillance and response capabilities (Hodcroft et al., 2021; Ye et al., 2021; Morel et al., 2021).

These shortcomings have spurred new research initiatives into phylogenetic inference methods capable of analyzing millions of samples. In particular, there has been significant recent progress in estimating and/or placing novel sequences onto very large phylogenies (Minh et al., 2020; Turakhia et al., 2021a; Aksamentov et al., 2021; Ye et al., 2022a,b). Accurate estimation of the underlying phylogeny has numerous downstream applications, including contact tracing (e.g., Lam-Hine et al., 2021; McBroome et al., 2022), surveillance (e.g., Abe and Arita, 2021; Klink et al., 2021), and improved understanding of pathogen biology (e.g. Majumdar and Sarkar, 2021; Turakhia et al., 2021b).

Another area of active research in phylogenetics, distinct from tree inference, is so-called *phylodynamics*, which seeks to understand how immunological, epidemiological, and evolutionary forces interact to shape viral phylogenies (Volz et al., 2013). Here, the quantity of interest is typically a low-dimensional parameter vector characterizing the underlying phylodynamic model, while the phylogeny itself is a nuisance parameter. Of particular interest for the current pandemic are methods that can estimate effective population size and reproduction number of the pathogen from viral genetic data (e.g. Zhou et al., 2020; Lai et al., 2020; Volz et al., 2021; Campbell et al., 2021). Compared to phylogeny estimation, less progress has been made on so-called “phylodynamic inference” at the pandemic scale. This absence motivates the present study.

Bayesian methods are often preferred for phylodynamic inference because there are usually many trees which explain the data equally well. Hence, downstream quantities of interest possess a potentially significant amount of “phylogenetic uncertainty” which is not reflected in frequentist point estimates. Unfortunately, Bayesian phylogenetic procedures inherently scale very poorly: the space of phylogenetic trees grows rapidly, and there are an astronomical number of possible trees to consider, even for relatively small samples. Consequently, on large problems, the workhorse algorithm of field, Markov chain Monte Carlo (MCMC), tends to either conservatively explore very limited regions of tree space, or liberally propose large moves that are often rejected (Whidden and Matsen IV, 2015; Zhang and Matsen IV, 2019).

Even before the pandemic, awareness of the scalability issues surrounding Bayesian phylogenetics was growing (Höhna and Drummond, 2012; Whidden and Matsen IV, 2015; Aberer et al., 2016; Dinh et al., 2017). As a scalable alternative to MCMC, variational inference (VI) has recently garnered some attention in phylogenetics. VI is a general method for sampling approximately from a posterior distribution using techniques from optimization (Jordan et al., 1999). Fourment et al. (2020) used VI to accelerate computation of the marginal likelihood of a fixed tree topology. Fourment and Darling (2019) used the probabilistic programming language STAN to perform variational inference of the Bayesian skyline model (Pybus et al., 2000). Both of the preceding methods only analyze a fixed tree topology, so they cannot account for phylogenetic uncertainty. Simultaneously, Zhang and Matsen IV (2018, 2019); Zhang (2020) have made progress on a full variational approach which includes optimization over the underlying topology. Although these innovations represent significant advances in terms of performance, they still cannot come close to exploiting all of the information contained in a pandemic-scale data set.

## 2 New Approaches

Inspired by these works, and responding to the need for better tooling to study the ongoing pandemic, we devised a method capable of providing accurate and calibrated estimates of the rates of transmission and recovery for COVID-19 using data from tens of thousands of viral genomes. Our approach unites several threads of research in phylogenetics and scalable Bayesian inference. We build on aforementioned advances in variational phylogenetic inference (Fourment and Darling, 2019; Zhang, 2020), as well as recent progress in phylodynamic modeling of infectious diseases (Stadler et al., 2013), Bayesian stochastic optimization (Hoffman et al., 2013), and differentiable programming (Bradbury et al., 2018). To achieve this level of scalability, our method makes several tradeoffs and approximations which are detailed below. Briefly, we adopt a divide-and-conquer strategy where distant subtrees of a very large phylogeny are assumed to evolve approximately independently, and we further assume that topological estimates of these subtrees are an accurate reflection of their distribution under the prior. We argue that these are reasonable approximations in the context of an massive, global phylogeny, and that their combined effect appears to be benign: the resulting estimates closely agree with the existing state of the art on simulated data, and exhibit a remarkable level of concordance with ground-truth estimates on real data, while taking just minutes to produce.

## 3 Results

In this section, we test our method on both simulated and real data, and compare it to the existing implementation of the birth-death skyline model in BEAST.

### 3.1 Simulation

First, we performed a simulation study to evaluate how well VBSKY approximates the posterior distribution compared to BEAST. We studied four different scenarios:

1. Constant: the effective reproductive number stays constant through time;
2. Decrease: there is a sharp drop in the effective reproductive number;
3. Increase: there is a sharp increase in the effective reproductive number; and
4. Zigzag: the effective reproductive number goes through a series of decreases and increases.

We simulated transmission trees using the `R` package TreeSim (Stadler, 2011) and generated sequences data along each tree using the program Seq-Gen (Rambaut and Grass, 1997).

Across all scenarios, the rate of becoming uninfectious, *δ* is held constant at *δ*(*t*) = 4 for all *t*. The sampling rate is also held constant at *s*(*t*) = 0.25. Only *R* is allowed to vary. Under the constant scenario, *R*(*t*) = 1.3 for all *t*. In the decrease scenario,

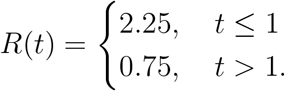

In the increase scenario,

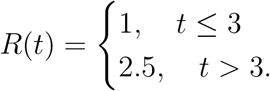

In the zigzag scenario,

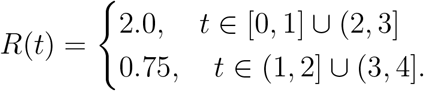

Each simulation was run for four time units, and ten trees were generated under each scenario. Because the sampling process is stochastic in this model, the size of the simulated tree varied from run to run. The minimum (maximum) number of samples in each under the constant, decrease, increase, and zigzag scenarios was 175 (1553), 117 (590), 124 (1075), and 161 (1852), respectively.

We compared the performance of our method to the current state-of-the-art for Bayesian phylogenetic analysis, BEAST (Bouckaert et al., 2019). BEAST allows for the birth-death skyline model to be used as a tree prior, facilitating direct comparison with VBSKY. Because BEAST uses MCMC to estimate the posterior, the number of sequences it can analyze is limited. Therefore, for each simulation, we randomly sampled 100 sequences for BEAST to analyze. We allowed BEAST to run long enough that the effective sample size exceeded 1000 for each evolutionary parameter. Since VBSKY is not limited by sample size, we analyzed all sequences in each simulation, as follows: We set the size of each random subsample to be *b* = 100 tips. The number of trees in the ensemble was set to be the smallest integer such that the number of trees multiplied by 100 was larger than the number of sampled sequences. Under this scheme, each sequence was sampled once on average.

The results of the simulation study are shown in Figures 1 and 2. Figure 1 displays the median of the medians and 95% equal-tailed credible intervals of the simulations under each scenario using BEAST to analyze the data. Figure 2 shows the same for VBSKY. Besides a few minor differences, the estimates given using VBSKY are similar to those given by BEAST; both BEAST and VBSKY adequately capture the true value of the effective reproductive number. The credible intervals given by BEAST are wider than those of VBSKY, and do a better job of covering the true model in some cases; we return to this point in Section 4. In the decrease scenario, VBSKY is better able to capture the larger value of *R* earlier in time, while BEAST appears to revert to the prior at times earlier than *t* = 0.5. Because VBSKY allows for more sequences to be analyzed, the method is able to detect transmission events further back in time.

**Figure 1:**
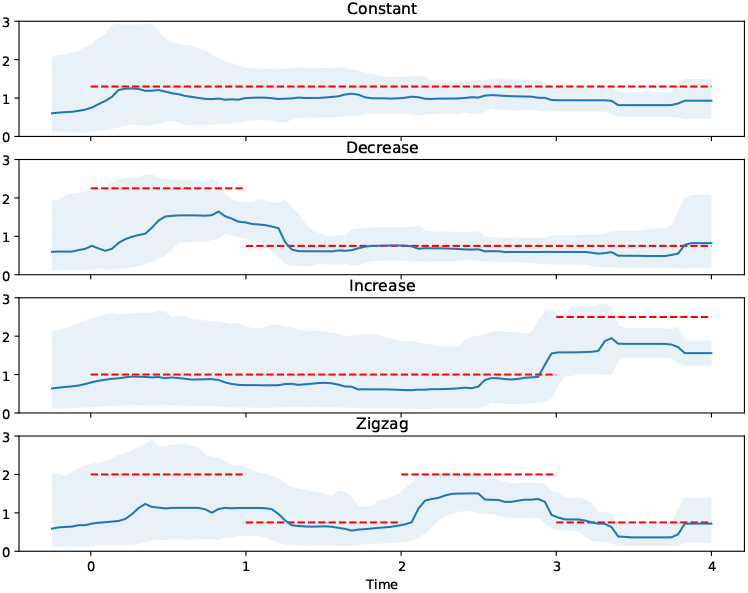
Median of the medians and the equal-tailed 95% credible intervals of the posteriors of the effective reproductive number over time of the 10 simulations for each scenario using BEAST. The dotted red line is the true effective reproductive number over time.

**Figure 2:**
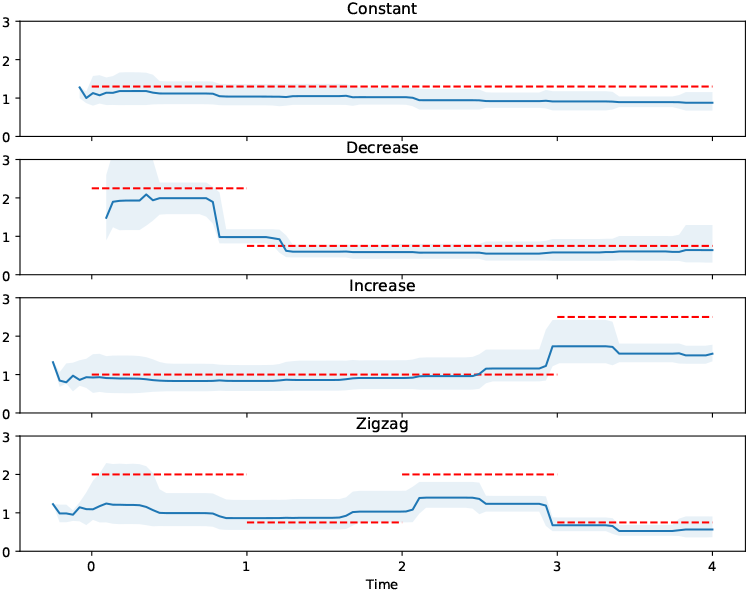
Median of the medians and the equal-tailed 95% credible intervals of the posteriors of the effective reproductive number over time of the 10 simulations for each scenario using VBSKY. The dotted red line is the true effective reproductive number over time.

Even though in some cases we analyzed hundreds more sequences using VBSKY than when we used BEAST, the run-time of VBSKY was 71.75 seconds on average for each simulation whereas BEAST took 20 minutes to perform 10,000,000 MCMC steps. The simulation results show that VBSKY is able to get comparable results as BEAST with a much shorter run-time, and in some cases like the decrease scenario, VBSKY can produce more accurate estimates than BEAST.

### 3.2 Analysis of the global pandemic

We tested our method on a large, serially-sampled COVID-19 dataset from the GISAID initiative (Elbe and Buckland-Merrett, 2017). At the time this analysis was performed, there were 6.5M SARS-CoV-2 sequences in the database. In addition to the raw nucleotide data, GISAID provides sample time and location information. The collection dates of the sequences range from January 3rd, 2020 to December 8th, 2021.

For our analysis, we chose to study the transmission of COVID-19 of Michigan, Florida, and the entire USA. It is important to study the epidemiology of COVID-19 at the sub-national level as many public health policies such as mask mandates, stay at home orders, vaccine distribution, and other social distancing measures are enforced at the state level. Policies or decisions made in one state may not be detected studying national data. Due to the differences in health policies across states and the reduced frequency of travel during the pandemic, we expect the incidence and prevalence of COVID-19 to vary from state to state. On the other hand, policies are sometimes made at the national level, and more recently travel especially around the holidays has become widespread, so understanding trends at a national level is equally vital.

After filtering the sequences by location, the number of sequences were 81,375, 34,978, and 1,280,563 for Florida, Michigan, and the USA respectively. We noticed that the number of confirmed cases increased or decreased based on the day of the week, likely because fewer cases are reported over the weekend. To correct for any inaccuracies in the sample time distribution, we set all sequences sampled in the same calendar week to have the same sample time. We used a fixed molecular clock model with substitution rate 1.12 × 10^−3^*/*bp*/*year which is the estimate given by the World Health Organization (WHO) (Koyama et al., 2020).

#### 3.2.1 Hyperparameter Tuning

Before proceeding to the analysis, we sought to better understand how the various tuning parameters of our method affected the results. VBSKY has two main tuning parameters that can be adjusted: the number of tips in each subsample (denoted *b* in the preceding section), and the number of subsamples of the overall dataset 𝒟 (denoted *S* in the preceding section). Increasing either enables us to analyze more sequences, but at the expense of additional computation time.

To understand the effect of the number of trees, we examined the posterior of the effective reproductive number and the sampling rate of Florida and the USA while fixing the number of tips and varying the number of trees. We set the number of tips to be 200 and examined the posterior for each number of trees in the set {10, 25, 50, 100, 150}. Patients with mild bouts of COVID-19 are generally not infectious after 10 days of symptom onset (Arons et al., 2020; Bullard et al., 2020). The rate of becoming uninfectious is the inverse of the number of infectious days. As one unit of time corresponds to one year, the estimated value for *δ* is given by 1*/*10 × 365 = 36.5. Using this, we fixed the uninfectious rate to be 36.5 to avoid nonidentifiability issues since we cannot estimate *R, δ*, and *s* simultaneously (Stadler, 2009; Louca and Pennell, 2020). For the GMRF smoothing prior, we chose a relatively uninformative hyperprior distribution with large variance for the parameters of the smoothing prior. In particular, we selected a gamma distribution distribution with parameters *a* = *b* = 0.001, giving a mean of 1 and variance of 1000. As a rough estimate of the sampling rate, we also chose the prior for *s* to be a Beta(0.02, 0.98) distribution with expectation 0.02, as the ratio of sampled sequences to the number of cumulative cases is around 0.02. The remaining priors are shown in the first line of Table 1.

**Table 1:** Prior Distributions used in Analyses.

| Analysis | $R$ | $s$ | $\tau_R$ | $\tau_s$ | $x_1$ |
| --- | --- | --- | --- | --- | --- |
| Uninformative Smoothing | $\text{LogN}(1,1)$ | $\text{Beta}(.02, .98)$ | $\text{Gamma}(.001, 0.001)$ | $\text{Gamma}(.001, 0.001)$ | $\text{LogN}(-1.2, 0.1)$ |
| Less Smoothing | $\text{LogN}(1,1)$ | $\text{Beta}(20, 980)$ | $\text{Gamma}(10, 100)$ | $\text{Gamma}(10, 100)$ | $\text{LogN}(-1.2, 0.1)$ |
| Biased Sampling | $\text{LogN}(1,1)$ | $\text{Beta}(20, 980)$ | $\text{Gamma}(.001, 0.001)$ | $\text{Gamma}(.001, 0.001)$ | - |
| Multistrain | $\text{LogN}(1,1)$ | $\text{Beta}(.02, .98)$ | $\text{Gamma}(10000, 0.01)$ | $\text{Gamma}(.001, 0.001)$ | $\text{LogN}(-1.2, 0.1)$ |

Figure 3 shows the posterior of *R* for both Florida and the USA when varying the number of trees. Figure 4 shows the posterior for *s*. The figures indicate a larger difference when the number of trees is 10 compared to any greater number of trees. The median and credible interval for *R* was much smaller and the median and credible interval for *s* was much larger closer to the present when the number of trees was 10. The credible intervals when the number of trees was 10 was also much wider. A closer inspection showed that this also seems to be the case when the number of trees is 25, albeit to a smaller degree. When we increased the number of trees to 50, this difference mostly disappeared.

**Figure 3:**
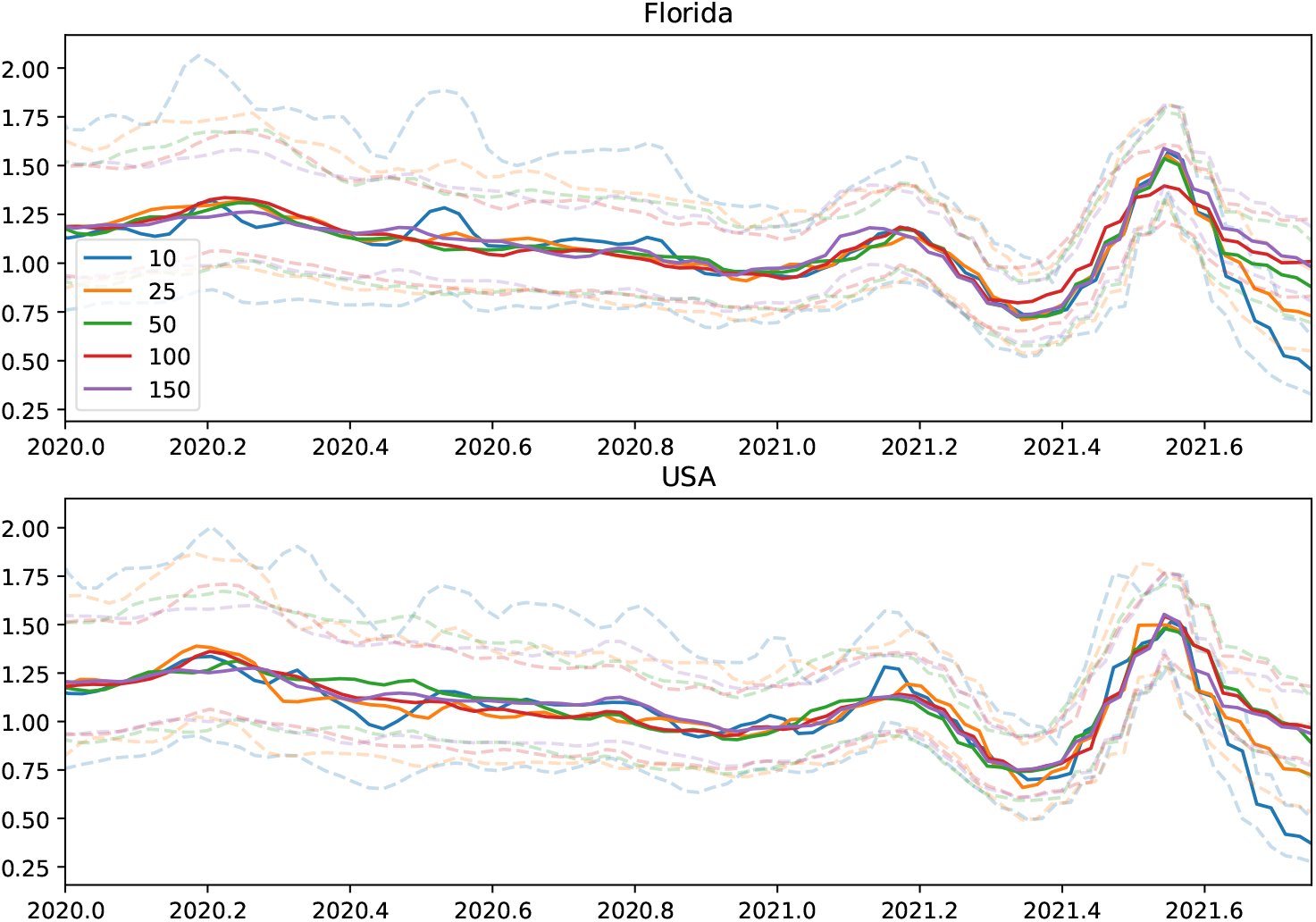
Posterior of R while varying the number of trees. Solid lines represent the median and the dotted lines represent the equal-tailed 95% credible intervals.

**Figure 4:**
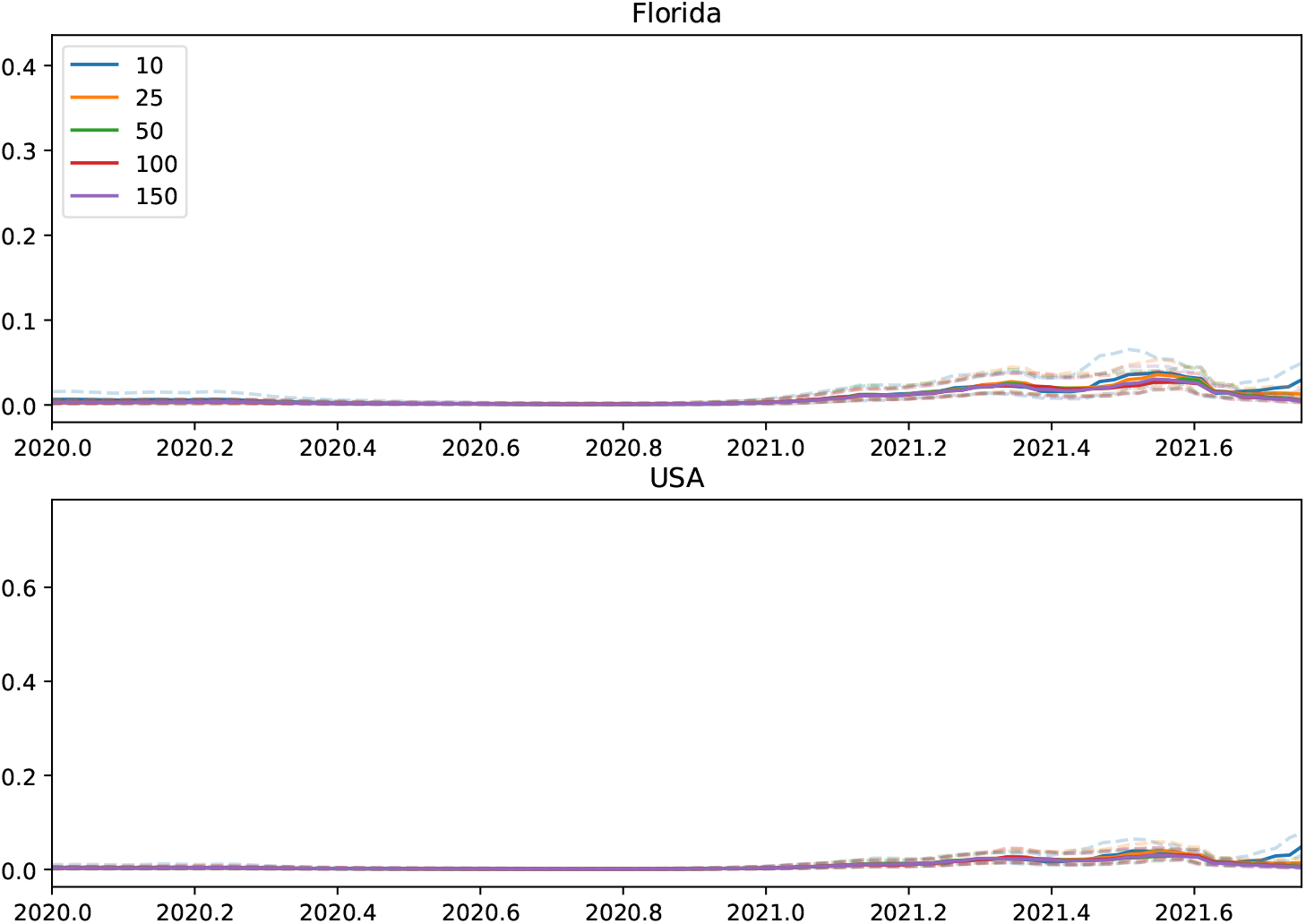
Posterior of ***s*** while varying the number of trees. Solid lines represent the median and the dotted lines represent the equal-tailed 95% credible intervals.

We performed a similar study to understand the effect of varying the number of tips. We fixed the number of trees as 50 and adjust the number of tips to values in the set {50, 100, 200, 400}, and examined the posteriors of *R* and *s* while holding *δ* fixed. Similar to above, varying the number of tips does not appear to have a large effect on the results. Using only 50 tips per tree resulted in a wider credible interval for Florida and the USA for both *R* and *s*. Figure 5 shows that using 50 tips also leads to flatter estimates for *R* further back in the past. This is likely the result of trees with fewer tips having fewer transmission events further back in the past which can be used to estimate *R*.

**Figure 5:**
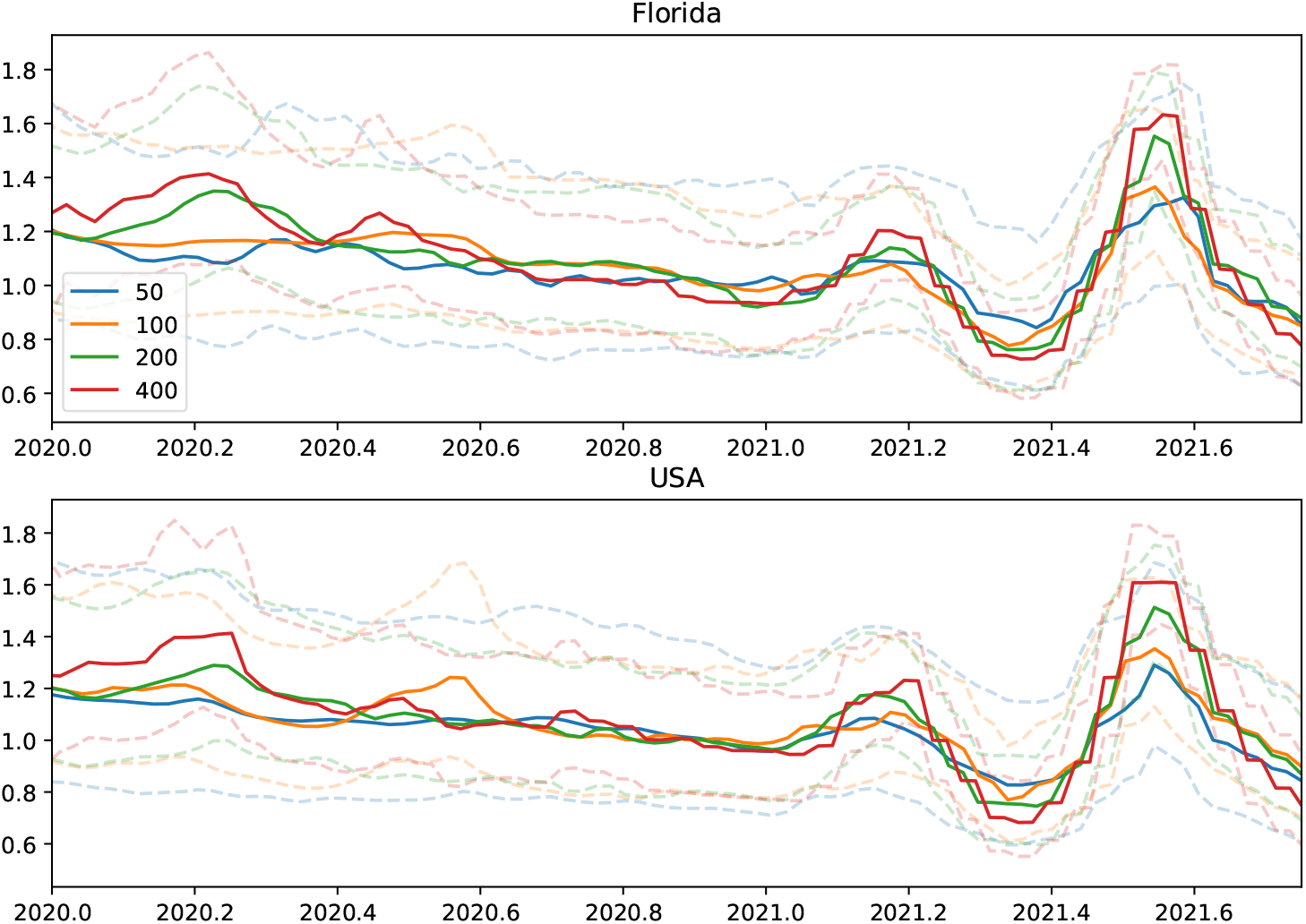
Posterior of R while varying the number of tips. Solid lines represent the median and the dotted lines represent the equal-tailed 95% credible intervals.

When comparing the posteriors when the number of tips is 100 or 200, only minor differences appeared. Using 200 tips did seem to lead to better detection of changes in *R* and *s* further back in the past. Looking at Figure 5, using 400 tips per tree led to a sharper decrease in *R* towards the present. Figure 6 shows that using 400 tips generally led to slightly larger estimates of *s* at all points in time.

**Figure 6:**
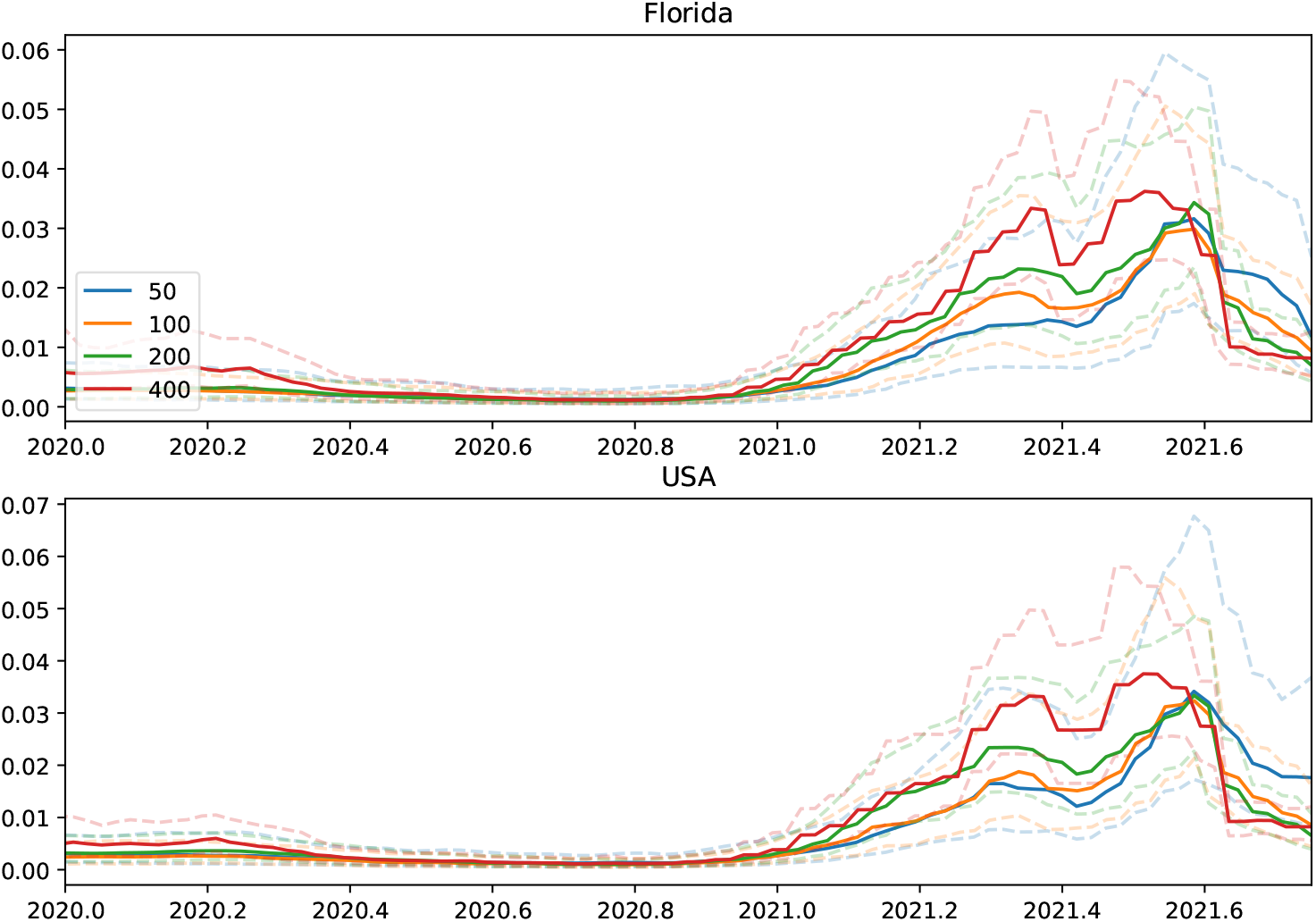
Posterior of s while varying the number of tips. Solid lines represent the median and the dotted lines represent the equal-tailed 95% credible intervals.

Overall, regardless of the number of tips or trees used, the posterior estimates of both *R* and *s* for both Florida and the USA are similar. However, increasing the number of trees decreases the variances in posterior estimates of *R* and *s*, and also results in more accurate estimates of both parameters towards the present. This improvement seems to plateau after increasing the number of trees to 50. Similarly, increasing the number of tips can increase the power to detect changes in *R* and *s* further back in the past, but using too many tips can lead to more erratic estimates of the parameters towards the present.

Keeping this in mind while also noting that increasing the number of trees and tips can incur large computational costs, using 50 trees with 200 tips leads to sharper estimates of the posterior without requiring excessive computation.

#### 3.2.2 Results

Based on the results from the previous section, we ran VBSKY with 50 subsamples of 200 sequences for a total of 10^4^ sequences. We estimated the epidemiological parameters for Florida, Michigan, and the overall USA. State-level results were compared to a “ground truth” estimator of the effective reproductive number which is derived from orthogonal (i.e. non-genetic) public health data sources (Shi et al., 2021). The prior and hyperprior settings for all of the scenarios described below are shown in Table 1.

We first analyzed the data using the same uninformative smoothing hyperpriors as in the hyperparameter study in the previous section (“Uninformative Smoothing” in Table 1). Figure 7 displays the posterior of *R* over time for each region for the uninformative smoothing analysis. For Florida (top panel), we see that the estimates for *R* over time produced by VBSKY matches the results using surveillance data in the recent past. However, earlier in the pandemic, VBSKY does not seem to be able to capture the rise and fall of *R* but instead provides a flat estimate of the parameter.

**Figure 7:**
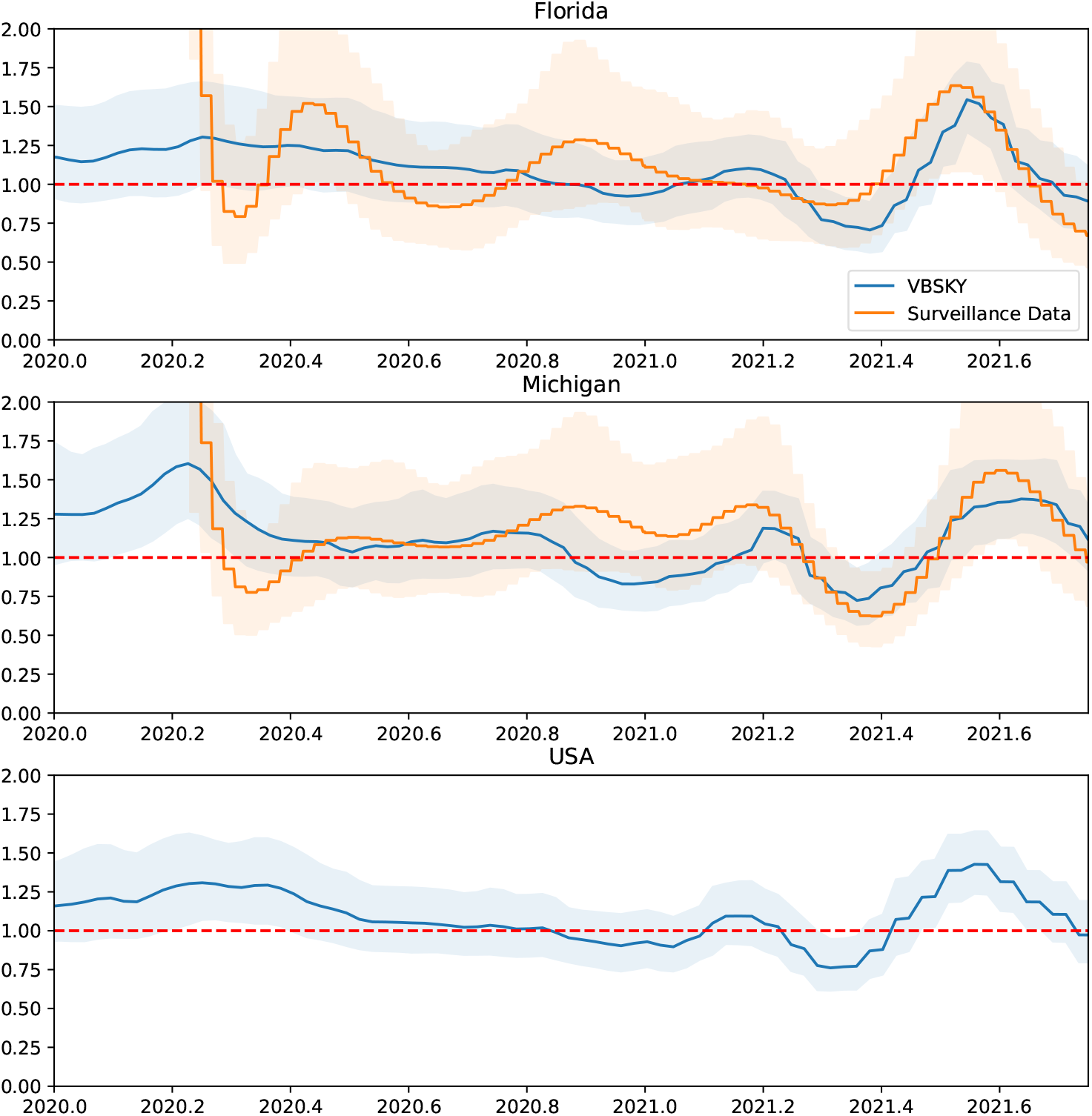
Posterior of *R* for Florida, Michigan, and the USA using an uninformative smoothing prior. VBSKY estimates are in blue. The orange estimates are derived from surveillance data. For each method the posterior median and equal-tailed 95% credible interval are shown. The dotted red line is *R* = 1.

In the middle panel (Michigan), we see the VBSKY posterior is very similar to the posterior given by the surveillance data method even looking further back in the past. Looking at the top panel (USA), similar to the results for Florida, the posterior for *R* is very flat further back in the past. Given that we have seen large rises and falls in the number of cases over time (Figure S2), it seems unlikely that the actual value of *R* is as flat as the method suggests.

One explanation for this performance discrepancy is that the prior may be oversmoothing the estimates of *R* further back in the past for some of the data sets. Figure S1 shows the distribution of sample times for Florida, Michigan, and the USA. Michigan has a larger proportion of sequences sampled early in the pandemic compared to either Florida or the overall USA. Oversmoothing may occur because a lack of samples further back in the past causes the prior to overwhelm the data.

To investigate this, we reran the analysis with stronger hyperpriors designed to reduce the overall amount of smoothness (“Less Smoothing” in Table 1). Figure 8 shows the posterior when we set the prior of the smoothing parameter to be a gamma distribution with *a* = 10 and *b* = 100, giving a mean of 0.1 and variance 0.001. Looking at the top panel (Florida) of Figure 8, we see that the posterior median of *R* for VBSKY is no longer flat and instead oscillates to better match the results using surveillance data. The bottom panel (USA) also shows the estimates for *R* for the entire USA are also no longer completely flat further back in the past. The middle panel (Michigan) shows that even with less smoothing, the results for VBSKY in Michigan match well with the surveillance data. When the sample time distribution is unbalanced, as with Florida and the USA, imposing less smoothing can help better capture the signal where the sampling may be more sparse. However, it also widens the credible intervals.

**Figure 8:**
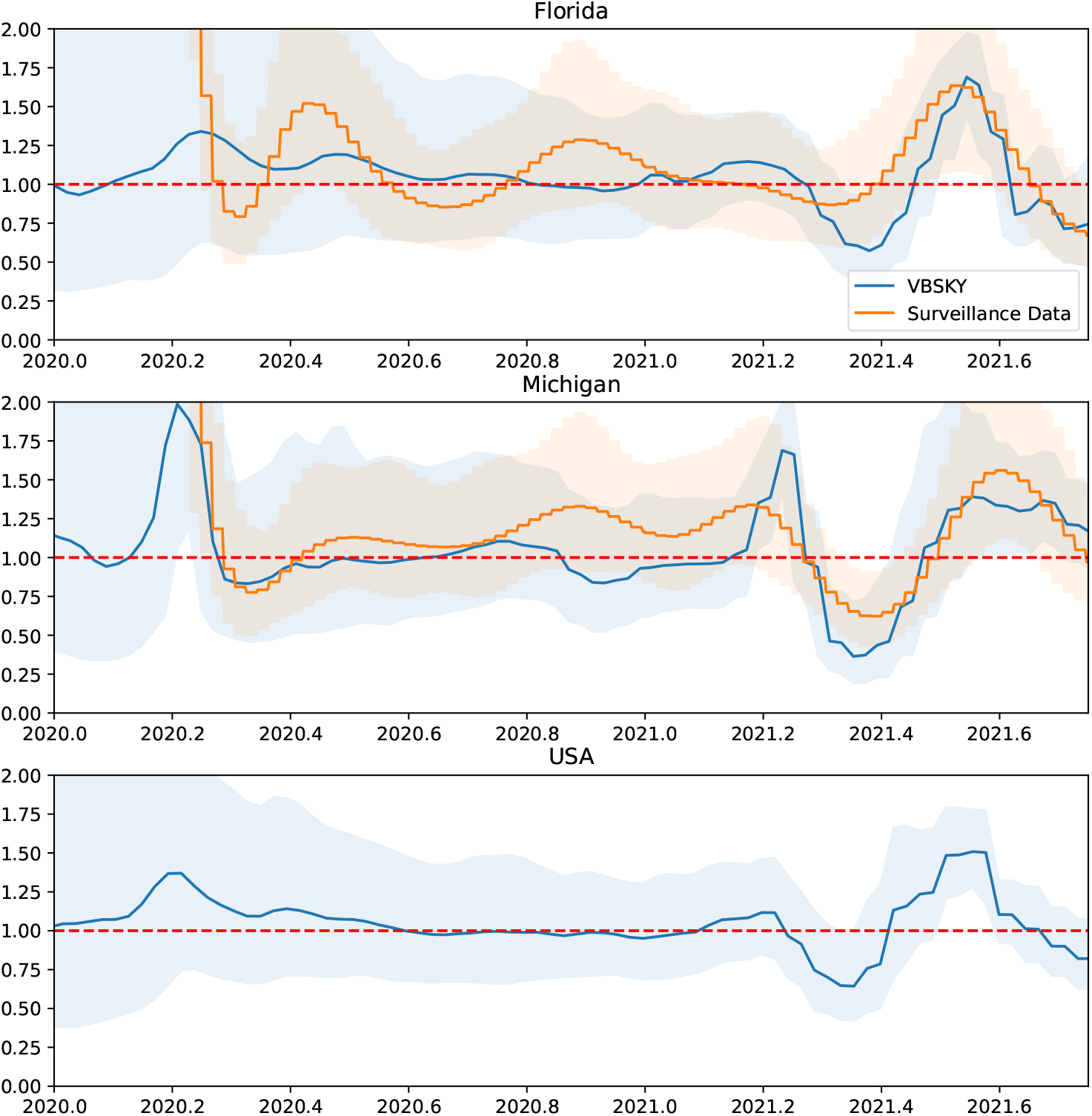
Posterior of *R* for Florida, Michigan, and the USA using less smoothing. VBSKY estimates are in blue. The orange estimates are derived from surveillance data. For each method the posterior median and equal-tailed 95% credible interval are shown. The dotted red line is *R* = 1.

In addition to decreasing the amount of smoothing, we explored the use of a biased sampling scheme to yield sharper estimates further back in the past. The algorithm described in Section 5 generates an ensemble of trees by sampling the data randomly without replacement. Hence, if most of the samples were collected in the recent past, most of the trees in the ensemble will have tips from near the present, making it difficult to estimate transmission events further back in time. To verify this, we split the data by the quarter in which the sequence was sampled, where the first quarter of each year was defined to be the first three months (January, February, March) of the year, and so on. Then, instead of randomly sampling to generate the ensemble of trees, for each tree the tips were restricted to only one quarter. We also enforced the number of trees per quarter to be approximately equal. One caveat is that this stratified sampling approach could bias the estimates of the sampling rate.

Figure 9 shows the results using the biased sample approach. For this final analysis we reverted some of the smoothing prior changes (“Biased Sampling” in Table 1). (Because of convergence issues encountered during model fitting, for this scenario we fixed the origin to 0.3 years prior to the earliest sample date; therefore, no prior on *x*_1_ is listed in the table.) There is a surprisingly close match between our model output and the ground-truth, which we reiterate was estimated using a completely different source of data. The estimates using the biased sampling approach improve the estimates of *R* further back in the past especially for Florida. Using less smoothing, VBSKY was able to capture the shape of estimates using surveillance data, but the biased sampling approach results in a much closer estimate of *R* further back in the past. The credible bands produced by VBSKY tend to be narrower, which could reflect either differences in the underlying data or violations of the modeling assumptions described in Section 5. Interestingly, both methods appear unable to reject the null hypothesis *R* = 1 except for very early in the pandemic (winter 2020) and very recently (spring-summer 2021). One drawback of the stratified sampling approach is that the estimates of *R* towards the present seem to be further away from the estimates using surveillance data. While using the biased sampling approach can improve estimates within time periods where sampling is sparse, it can also bias the estimates where sparse sampling is not an issue.

**Figure 9:**
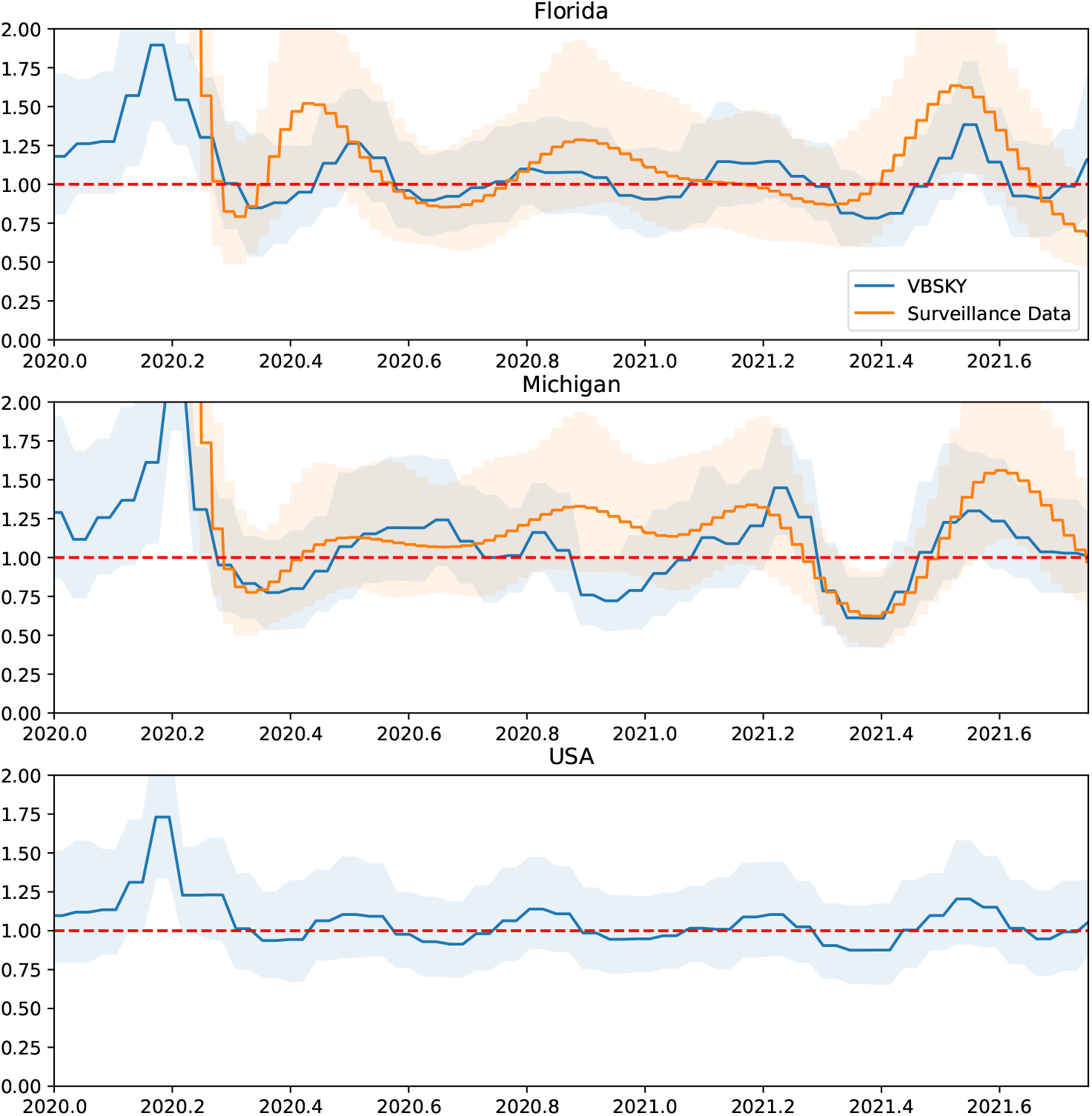
Posterior of *R* for Florida, Michigan, and the USA using biased sampling and a strong prior on *s*. VBSKY estimates are in blue. The orange estimates are derived from surveillance data. For each method the posterior median and equal-tailed 95% credible interval are shown. The dotted red line is *R* = 1.

In this section we focused on estimating the effective reproduction number *R*. A parallel set of estimates for the sampling fraction *s* are shown in Figures S3–S5.

#### 3.2.3 Comparison to BEAST

We ran BEAST on the same data set as in the previous section. BEAST was incapable of analyzing the same number of samples as VBSKY, so to facilitate comparison, we limited the number of sequences we analyzed with BEAST. Both the sample size and the sampling scheme can affect the results of the analysis as well as the mixing time, so we compared how BEAST performed with different combinations of sample sizes and sampling schemes. We ran BEAST with both 100 and 500 sequences. For each sample size, we sampled the most recent sequences by date (contemporary sampling), and we also sampled uniformly at random without any regard to the sample time (random sampling). The XML configuration files we used to run BEAST are included in the supplementary data.

Even after greatly reducing the number of sequences analyzed, accurately sampling from the posterior may still take longer than using VBSKY. We performed both a “short” run for BEAST, where the MCMC sampler is only allowed to run for as long as it took VBSKY to analyze the full data, as well as a “long” run where BEAST was allowed to perform 100 MCMC million iterations, or run for 24 hours, whichever was shorter.

The estimates of the effective reproductive number of the short run for Florida, Michigan, and the USA are displayed in Figures 10, S6, and S7 respectively. The estimates for the long runs are shown in Figures 11, S8, and S9.

**Figure 10:**
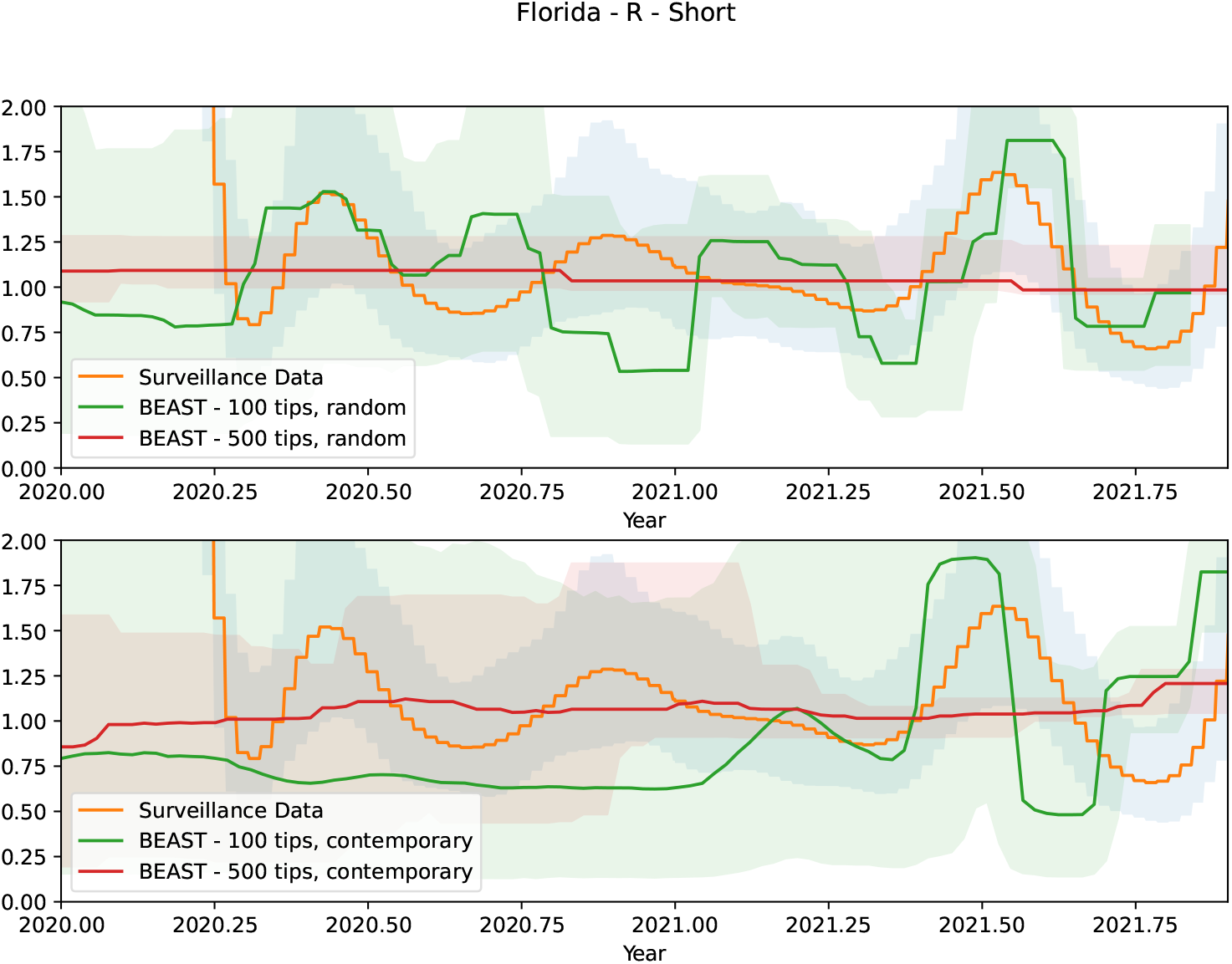
The posterior median and equal-tailed 95% credible interval of *R* for Florida given by BEAST. The top panel contains randomly sampled data, while the bottom contains the most recent available samples. The sampler was allowed to run as long as it VBSKY to analyze the Florida data. This is referred to as the short run in the text.

**Figure 11:**
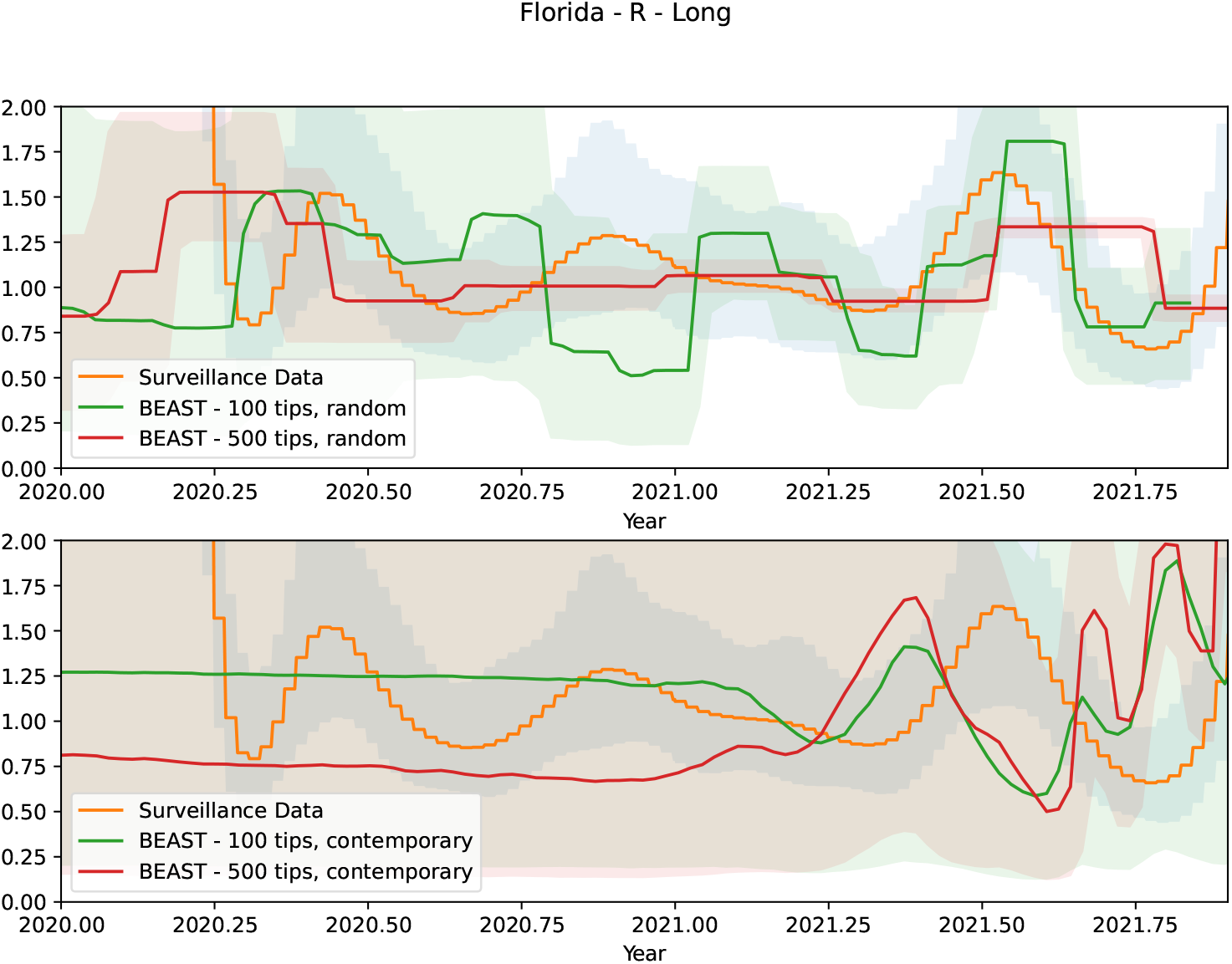
The posterior median and equal-tailed 95% credible interval of *R* for Florida given by BEAST. The sampler was allowed to run for 100 million steps or 24 hours to analyze the data. This is referred to as the long run in the text.

For the short runs, depending on the number of samples and the sampling scheme, the results varied widely. Under a short time constraint, the posteriors using 500 tips and both sampling schemes for Florida, 500 tips and recent sampling for Michigan and 500 tips and recent sampling for the USA were mostly flat centered close to 1. The posteriors did not reflect the rise and fall in *R* that is exhibited in both the surveillance data and VBSKY estimates. In most cases, BEAST is unable to capture any signal further back in the past, and the posterior provided by BEAST does not track the estimates provided by the surveillance data as well as VBSKY.

In the long runs, the issue of completely flat posteriors when using 500 tips mostly disappeared. However, BEAST is only capable of producing comparable results to VBSKY and the surveillance method when analyzing 100 tips sampled uniformly at random, presumably because mixing occurred more rapidly in the time allotted. The long runs also illustrate that uniform random sampling performs better than most-recent sampling when running BEAST. This indicates that having samples throughout time may help infer more transmission events further back in the past rather than having only contemporary sequences. The discrepancy between using 100 tips and 500 tips exists only when the sampling scheme is random. When using contemporary sequences, BEAST is able to complete 100 million iterations. But when random sampling is used, because the MCMC sampler mixes more slowly, BEAST was unable to complete 100 million MCMC moves within 24 hours.

In summary, BEAST performed fairly well when we randomly sample 100 tips, though there was considerable variation between data sets and scenarios. The main difference between VBSKY and BEAST is that the latter was usually unable to capture signal far back in the past. Analyzing more sequences could help, but the computational difficulties that would ensue imply that it is not practical to completely resolve this issue if time is a constraint. Overall, our results indicate that efficiently analyzing thousands of sequences, even using an approximate inference method, generally leads to a sharper posterior which is closer to the ground truth.

#### 3.2.4 Strain Analysis

As a supplement to our main analysis, we further investigated the history of different COVID-19 variants. Using GISAID-annotated variant information, we split our data set of Florida, Michigan, and USA sequences into smaller data sets specific to the Alpha and Delta variant and fit our model to each variant.^1^ Except for a minor adjustment to the prior on the origin time, we used all the same hyperparameters and priors as in the preceding section. For the GMRF smoothing prior, we chose a hyperior for *τ*_*R*_ to have large expectation to increase smoothing.

The results of our analysis are shown in Figure 12 for *R* and Figure S10 for *s*. The Alpha variant of COVID-19, also known as lineage B.1.1.7, originated in England and was first reported in the USA in early 2021. Using surveillance data, Volz et al. (2021) showed that at the time, the Alpha variant had a transmission advantage over other variants, which is why it came to dominate in the USA in early 2021. There are no samples for the Alpha variant beyond summer 2021, so the estimates for Alpha are truncated at various points during that period depending on the region considered. As shown in Figure S2, the number of cases in Michigan, Florida, and the USA all dropped after the first third of the year, corresponding to a decrease in *R* below one for the Alpha variant. At the same time, the Delta variant was rising in prevalence, such that *R* is estimated greater than one in all cases until about the third quarter of 2021. Analysis of the sampling fraction over time (Figure S10) also shows some interesting trends, for example sampling of the Delta variant in Michigan seems to have been extremely low compared to other areas and strains. Finally, we also explored using other hyperparameter settings to analyze these data, but found that they produced suboptimal results. In particular, without additional smoothing, our model unrealistically estimated that *R* increased for the Alpha variant throughout the second quarter of 2021, although the credible intervals generally place substantial posterior probability on the event *R <* 1 (Figures S11 and S12). We noticed that for the Alpha variant, the number of available samples drops severely near the point of truncation. The absence of data would lead to the prior dominating the posterior samples of *R*. By increasing smoothing, we were able to circumvent this issue.

**Figure 12:**
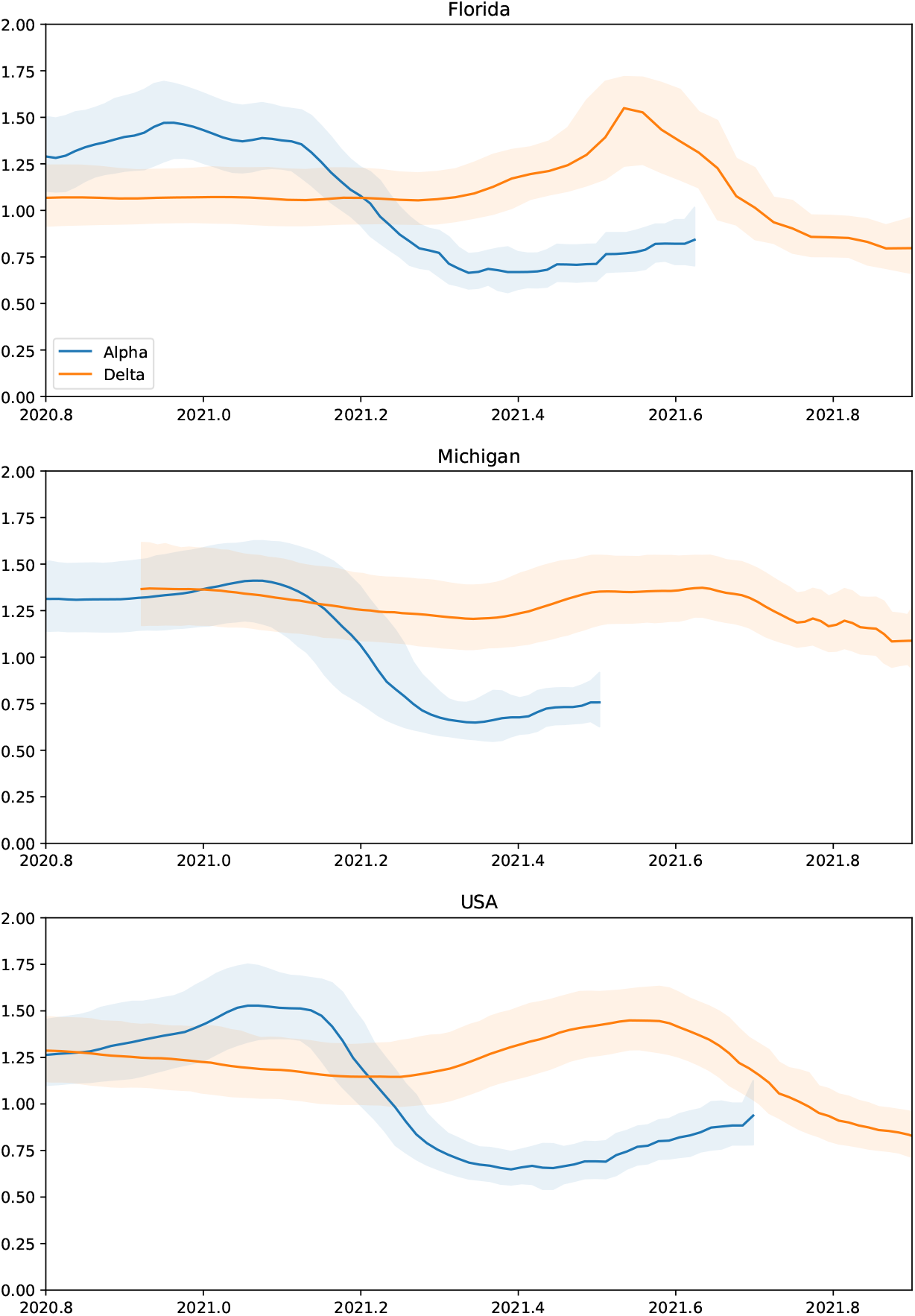
The posterior median and equal-tailed 95% credible interval of *R* for the Alpha and Delta variants.

## 4 Discussion

In this paper, we presented the variational Bayesian skyline, a method designed to infer evolutionary models from large phylogenetic datasets. Our method works by fitting a variational Bayesian posterior distribution to a certain approximation of the phylogenetic birth-death model. We showed that, under some simplifying heuristic assumptions, it can be used for posterior inference of epidemiologically relevant quantities such as the effective reproduction number and sampling fraction. We demonstrated that our estimates adhere reasonably closely to alternative approaches such as MCMC, while being significantly faster and therefore able to incorporate large numbers of observations. On real data, we showed how our model corroborates public health surveillance estimates, and could work to fill in the gaps when such data are unavailable.

One shortcoming of our model is that it tends to be overconfident, in the sense that it produces credible intervals which are narrower compared to other methods, and not as well calibrated in simulations. Generally, it is preferable for a method to overcover since this is inferentially more conservative. We believe this behavior is attributable to the heuristics that underlie our approach: since they ignore certain forms of dependence in the data, they create the illusion of a larger sample size than actually exists. We suggest that the credible intervals produce by our method are best interpreted relatively, as showcasing portions of time where the estimates are especially sharp or loose.

Our method could be extended in several ways. Currently, it estimates the tree topology and the continuous variables separately, relying on a distance-based method infer the topology. While faster, distance-based methods are less accurate than likelihood-based methods for tree reconstruction (Kuhner and Felsenstein, 1994). Our method could be potentially extended to unify the estimating procedure for tree topologies and other variables under one variational framework allowing (Zhang and Matsen IV, 2019). We also take random subsamples of data to accelerate our inference. However, the subsampling approach we adopt is very naive, and future work could include developing an improved way strategy for subsampling in phylogenetic problems.

The variational inference scheme we used makes a standard but highly simplified mean-field assumption about the dependence structure of the variational approximating family. We also experimented with other, recent approaches such as normalizing flows (Rezende and Mohamed, 2015), but observed that, consistent with earlier findings (Fourment and Darling, 2019), they did not measurably improve the results and occasionally caused the algorithm to fail to converge. If our approach is adapted to more complex problems, it could be advantageous to revisit this modeling choice.

Currently, our method is restricted to using a strict molecular clock model. Additionally, the substitution models in our method do not currently allow for rate heterogeneity across sites. Allowing for more flexible and complex substitution and clock models could aid in the application of our method to other data sets that evolve differently than COVID-19, when the time scale of the epidemic is much larger.

## 5 Materials and Methods

In this section, we derive our method, which we call variational Bayesian skyline (VBSKY). As the name suggests, VBSKY descends from a lineage of earlier methods designed to infer evolutionary rate parameters from phylogenetic data (Pybus et al., 2000; Drummond et al., 2005; Minin et al., 2008; Gill et al., 2013). Our running example will be inferring the epidemiological history of the COVID-19 pandemic, but the method applies generally to any evolving system that is aptly modeled using a phylogenetic birth-death or coalescent process and approximately meets the assumptions described below.

### 5.1 Notation and model

The data consists of a matrix of aligned sequences **𝒟** = {*A, C, G, T, N*}^*n*×*L*^, where *n* is the number of viral sequences and *L* is the number of sites, and a vector of times when each sample was collected ***y*** = (*y*_1_,…, *y*_*n*_) where *y*_1_ ≤ · · · ≤ *y*_*n*_. Row *j* of **𝒟** corresponds to a sequenced viral genome collected from an infected host at time *y*_*j*_. Subsamples of *rows* of **𝒟** are denoted by 𝒟_*i*_ ∈ {*A, C, G, T, N*}^*b*×*L*^, with corresponding sample times 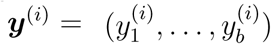, where *b* is the size of the subsample. We occasionally abuse notation and write 𝒟_*i*_ ⊂ **𝒟** to denote a subsample, and |𝒟| to denote the number of samples contained in a dataset (so e.g. |𝒟_*i*_| = *b* above). Phylogenetic trees are denoted by 𝒯 = (𝒯 ^topo^, 𝒯 ^br^), which we decompose into a discrete topological component and continuous branch length component. Given *n* sampled taxa, the topological component 𝒯 ^topo^ lives in the space of rooted, labeled bifurcating trees on *n* leaves, and the branch length component lives in the non-negative orthant 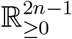 and gives the length of each edge of the tree (including an edge from crown to origin).

The data are assumed to be generated according to a phylogenetic birth-death skyline model (Nee et al., 1994; Morlon et al., 2011). In this model, samples are related by an unobserved “transmission tree” that records every infection event that occurred during the pandemic. Leaf nodes in the transmission tree represent sampling events, and internal nodes represent events where the virus was transmitted from one host to another. Edges denote periods during which the virus evolved within a particular host, with the length proportional to the amount of evolutionary time that elapsed between the parent and child nodes. The distribution of the infection tree depends on three fundamental parameters, usually denoted *μ*(*t*), *λ*(*t*), and *ρ*, which are respectively the time-varying per-capita rates at which extant lineages in the phylogeny go extinct and speciate, and the fraction of the extant population that was sampled at the present.

Further generalizations (Stadler et al., 2013) incorporate both random and deterministic sampling across time, and it was also shown how phylogenetic BD model can be used for parameter estimation in the susceptible-infected-recovered model (Kermack and McKendrick, 1927) that forms the foundation of quantitative epidemiology. Let *ψ*(*t*) denote the rate at which each extant lineage is sampled in the phylogeny. (Henceforth we suppress dependence on time, but all parameters are allowed to be time-varying.) If we assume that sampling is tantamount to recovery (a valid assumption when positive testing leads to quarantine, as is generally the case during the current pandemic), then the overall rate of becoming uninfectious is *δ* = *μ* + *ψ*; the average time to recovery is 1*/δ*; the sampling proportion is *s* = *ψ/δ*; and the effective reproduction number is *R* = *λ/δ*. Using prior knowledge, it is also common to specify an origin time *t*_0_ when the pandemic began.

Let *ζ* = (*R, δ, s, t*_0_) denote the vector of epidemiological parameters of interest. The hyperprior on *ζ* is denoted *π*(*ζ*). The latent transmission tree describing the shared evolutionary history of all of the sampled pathogens is denoted by **𝒯** = (**𝒯** ^topo^, **𝒯** ^br^). We assume a simple “strict clock” model, with known rates of substitution, so that no additional parameters are needed to complete the evolutionary model.

We desire to sample from the posterior distribution of *ζ* given the phylogenetic dataset **𝒟**. Let *p*(**𝒯** | *ζ*) denote the likelihood of the transmission tree given the evolutionary model. An expression for *p*(**𝒯** | *ζ*) can be found in Stadler et al. (2013, Theorem 1), and is reproduced in Appendix S-1 for completeness. The data depend on *ζ* only through **𝒯**, so that *p*(**𝒟** | **𝒯**, *ζ*) = *p*(**𝒟** | **𝒯**). Here *p*(**𝒟** | **𝒯**) denotes the “phylogenetic likelihood”, which can be efficiently evaluated using the pruning algorithm (Felsenstein, 1981). Putting everything together, the posterior distribution over the unobserved model parameters is

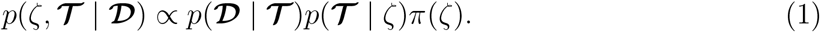

### 5.2 Scalable inference

The constant of proportionality in (1) is *p*(**𝒟**), the marginal likelihood after integrating out all (hyper)parameters and the unobserved tree **𝒯**. In large phylogenetic data sets, exact evaluation of the marginal likelihood is impossible due to the need to enumerate all possible trees, a set whose cardinality explodes in the number of taxa (Alfaro and Holder, 2006). In practice, methods such as Markov chain Monte Carlo (e.g., Drummond and Rambaut, 2007) which do not require evaluating *p*(**𝒟**) are utilized.

Since current phylogenetic MCMC algorithms cannot scale up to pandemic-sized datasets, we propose to modify the inference problem (1) using a few heuristics in order to make progress. Let 𝒟_1_, 𝒟_2_,…, 𝒟_*S*_ ⊂ **𝒟** be subsamples of *b*_1_,…, *b*_*S*_ rows from the full dataset. If the subsamples are temporally and geographically separated, and *b*_*i*_ ≪ *n*, then it is reasonable to suppose that these subsamples are approximately independent conditional on the underlying evolutionary model.

#### Heuristic 1.

*In a very large phylogenetic dataset* **𝒟**, *small subsets* 𝒟_1_, 𝒟_2_ ⊂ 𝒟 *with* |𝒟_1_|, |𝒟_2_| ≪ |**𝒟**| *that are sufficiently separated in space and/or time are approximately independent: p*(𝒟_1_, 𝒟_2_ | *ζ*) ≈ *p*(𝒟_1_ | *ζ*)*p*(𝒟_2_ | *ζ*).

True independence holds, for example, when the clades corresponding to 𝒟_1_, 𝒟_2_ are so distant that a reversible substitution process reaches stationarity on the edge connecting them. While we do not expect this to occur in real data, it seems like a reasonable approximation for studying distant subclades in a large, dense phylogeny which are evolving under a common evolutionary model. An example of the subsampling scheme we have in mind is when **𝒟** = “all of the samples collected in Florida” (*n* ≈ 81, 000), 𝒟_1_ = “all of the samples collected in Florida during June, 2020” (*b*_1_ ≈ 300), and 𝒟_2_ = “all of the samples collected in Florida during June, 2021” (*b*_2_ ≈ 5, 100).

Though incorrect, Heuristic 1 furnishes us with a useful formalism for performing large-scale inference, as we now demonstrate. Using the heuristic, we can approximate the posterior distribution (1) as

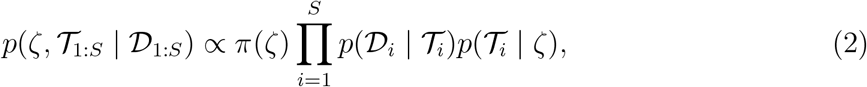

where we used the array notation 𝒯_1:*S*_ ≡ (𝒯 _1_,…, 𝒯_*S*_) to streamline the presentation. Sampling from (2) is easier than sampling from the full posterior (1) since it decomposes into independent subproblems, and each subtree 𝒯_*i*_ is much smaller than the global phylogeny 𝒯. However, the normalizing constant in (2) remains intractable even for small trees, so naäve sampling would still require expensive MCMC algorithms.

To work around this, we start by rewriting the last term in (2) as

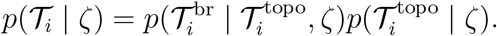

As noted in the introduction, the primary difficulty in Bayesian phylogenetic inference is navigating regions of topological tree space that have high posterior probability. If we could efficiently sample 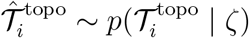, then the approximate posterior

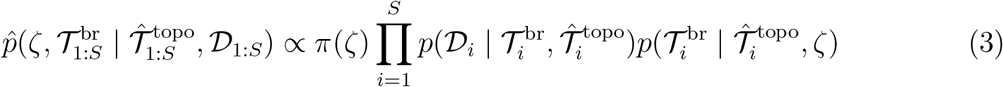

would have the property that

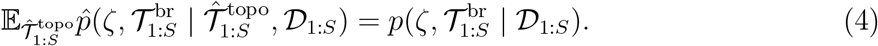

This leads to our second heuristic.

#### Heuristic 2.

*Fitted tree topologies* 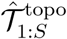 *obtained from subsets* 𝒟_1_,…, 𝒟_*m*_ *pairwise satisfying Heuristic 1 are independent and approximately distributed as p*(𝒯^topo^ | *ζ*).

By “fitted trees” we mean trees estimated using any method, including fast heuristic algorithms such as UPGMA, or its extension to serially-sampled time trees (sUPGMA; Drummond and Rodrigo, 2000); maximum likelihood; or simply extracting subtrees from a high-quality, pre-computed reference phylogeny (e.g., Lanfear, 2020). The heuristic can fail in various ways: in reality, tree reconstruction algorithms do not necessarily target the correct/any evolutionary prior, and there could be dependence between different trees if they are jointly estimated as part of a larger phylogeny. Also, our current implementation uses the data twice, once to estimate each tree, and again during model fitting to evaluate its phylogenetic likelihood. The tree inference procedure we used to analyze data in this paper is described more fully in the supplement (Section S-2). Note that we only utilize the *topological* information from these procedures; we still perform posterior inference over the branch lengths 𝒯^br^ as detailed below.

Setting these caveats aside, the point of Heuristic 2 is to endow our posterior estimates with some measure of phylogenetic uncertainty, without resorting to full-blown MCMC in tree space. By (4), the approximate likelihood (3) is unbiased for 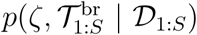, and the latter quantity correctly accounts for phylogenetic variance in the posterior. However, since (3) conditions on 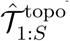, all of the remaining parameters to be sampled are continuous, and the problem becomes much easier.

Finally, we point out that our method is not capable generating useful samples from the posterior distribution *p*(**𝒯** | **𝒟**), that is of the overall transmission tree given the original dataset **𝒟**. But, as noted above, in skyline-type models the main object of interest is the evolutionary posterior *p*(*ζ* | **𝒟**). In Section 3, we demonstrate that the heuristic, subsampling-based approach developed here yields a fairly sharp posterior on *ζ*, while still utilizing a large amount of information from **𝒟**.

#### 5.2.1 Stochastic variational inference

Since (3) is a distribution over continuous, real-valued parameters, it is amenable to variational inference (Jordan et al., 1999). As noted in the introduction, variational Bayesian phylogenetic inference has previously been studied by Zhang and Matsen IV (2019); Zhang (2020) and Fourment and Darling (2019). Our approach is most related to the latter since we do not optimize over the topological parameters of our model in any way. Because we are operating in a different data regime than either of these two pre-pandemic papers, we further incorporated recent advances in large-scale Bayesian inference in order to improve the performance of our method.

Given a Bayesian inference problem consisting of data ***x*** and model parameters ***z***, traditional VI seeks to minimize the Kullback-Leibler (KL) divergence between the true posterior of interest and family of tractable approximating distributions 𝒬:

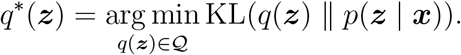

We cannot carry out this minimization as the KL divergence still requires evaluating the intractable quantity *p*(***x***). However,

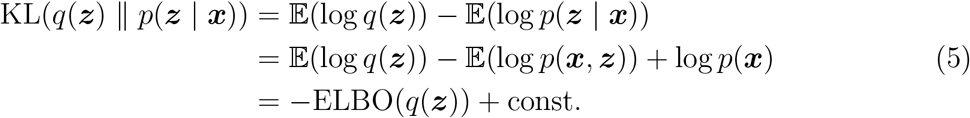

where the expectations are with respect to the variational distribution *q*, and

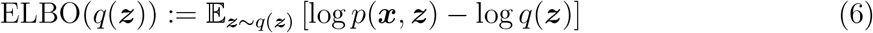

is known as the evidence lower bound. Hence, minimizing the divergence between the true and variational posterior distributions is equivalent to maximizing the ELBO.

For VI involving complex (non-exponential family) likelihoods, the ELBO is generally approximated by replacing the first term in (6) by a Monte Carlo estimate:

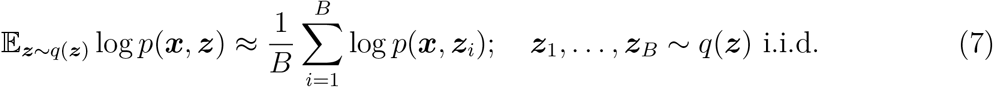

where *B* = 1 is a common choice. Each evaluation of the complete likelihood log *p*(***x, z***) requires a full pass over the data, which can be prohibitive when the data are large. Stochastic variational inference (SVI; Hoffman et al., 2013) addresses this problem through stochastic optimization. Many Bayesian models naturally factorize into a set of shared, global hidden variables, and sets of local hidden variables which are specific to each observation. Each observation is conditionally independent of all others given its local parameters. Hoffman et al. show how models of this form are well suited to stochastic gradient descent. Specifically, they derive an unbiased gradient estimator of the ELBO (6) which operates on a single, randomly sampled data point at each iteration. The algorithm tends to make better progress in early stages when the variational approximation to the shared global parameters is still quite inaccurate (Hoffman et al., 2013).

By design, the model we derived above is suited to SVI. In equation (3), the evolutionary parameters *ζ* are shared among all datasets, while the branch length parameters 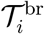 are specific to the *i*th dataset 𝒟_*i*_. We therefore refer to *ζ* as the global parameter, and the vectors of dataset-specific branch lengths 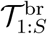 as local parameters. Our algorithm proceeds by iteratively sampling a single dataset 𝒟_*i*_ and taking a noisy (but unbiased) gradient step. Note that, because our model is not in the exponential family, we cannot employ the elegant coordinate-ascent scheme originally derived by Hoffman et al.. Instead, we numerically optimize the ELBO using differentiable programming (see below).

#### 5.2.2 Model parameterization

It remains to specify our model parameterization and the class of distributions 𝒬 that are used to approximate the posterior. Recall from Section 5.1 that the global parameter *ζ* includes the effective reproduction number *R*(*t*), rate of becoming uninfectious *δ*(*t*), and sampling fraction *s*(*t*). We follow earlier work (Gill et al., 2013) in assuming that these rate functions are piecewise constant over time, with changepoints whose location and number are fixed *a priori*. The changepoints are denoted **t** = (*t*_1_,…, *t*_*m*_) satisfying 0 = *t*_0_ *< t*_1_ *<* · · · *< t*_*m*_ *< t*_*m*+1_ = ∞. Thus,

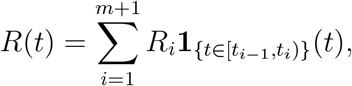

where the transmission rates in each time interval are denoted 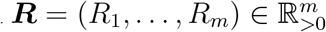. The rate of becoming uninfectious and sampling fraction are similarly denoted by 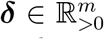 and ***s*** ∈ [0, 1]^*m*^, respectively. Finally, a Gaussian Markov random field (GMRF) smoothing prior is used to penalize consecutive differences in the log rates (Minin et al., 2008). To account for the fact that each rate parameter may have varying degrees of smoothness and also could be on different scales, each rate parameter has a corresponding precision hyperparameter *τ*_*R*_, *τ*_*δ*_, and *τ*_*s*_.

An extension of the BDSKY model allows for additional sampling efforts at each time *t*_*k*_. Infected individuals are sampled with probability *ρ*_*k*_ at time *t*_*k*_. When all sequences are sampled serially without the added sampling effort, *ρ*_*k*_ = 0 for 1 ≤ *k* ≤ *m*. When all sequences are sampled contemporaneously, ***ψ*** = **0**, *ρ*_*k*_ = 0 for 1 ≤ *k* ≤ *m* − 1, and *ρ*_*m*_ *>* 0. For our work, we only consider cases where *ρ*_*k*_ = 0 for 1 ≤ *k* ≤ *m* − 1. We define *b*_*s*_ as the number of sequences sampled serially, and *b*_*m*_ to be the number of sequences sampled at time *t*_*m*_. In other words, *b*_*m*_ is the number of contemporaneously sampled sequences at time *t*_*m*_. Note that *b* = *b*_*m*_ + *b*_*s*_. The sample times of the *b*_*s*_ serially sampled sequences are denoted by 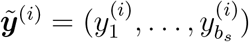. Because the sequences sampled at *t*_*m*_ have the largest sample time, 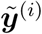 is just a truncated version of ***y***^(*i*)^. When all sequences are sampled serially, 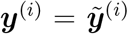. To conserve notation, from this point onward, we will use ***y***^(*i*)^ to refer to 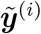.

The final remaining global parameter is the epidemic origin time *t*_0_. In order for the model to be well defined, this must occur earlier than the earliest sampling time in any of the *S* subsamples. Therefore, we set *t*_0_ + *x*_1_ = *y*_min_, where *y*_min_ is the earliest sampling time across all subsamples, and place a prior on *x*_1_ *>* 0 as detailed below.

Given the sampling times and estimated tree topology 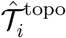, we can identify each local parameter 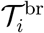 with a vector 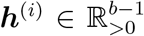 giving the height of each internal node when enumerated in preorder. Hence the height of the root node is 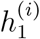. We follow the parameterizations set forth by Fourment and Darling (2019). In order for a sampled tree to be valid, we must have 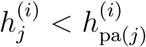 for every *j*. Here pa(*j*) denotes the parent node of node *j*. This constraint can be met by setting the height of internal node *j* as 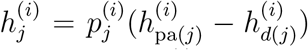 where *d*(*j*) is the earliest sampled tip from the set of descendants of *j* and 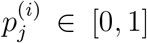. Finally, let 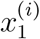 denote the distance of the root node from the origin measured forward in time. We must have 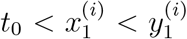 since the root node of 𝒯_*i*_ has to be between the origin and the earliest sample time. Therefore we set 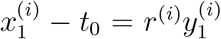 for some *r*^(*i*)^ ∈ [0, 1], and calculate the root height 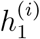 from it. Under this parameterization, the set of local variables 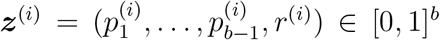 is a set of proportions, with transformations to switch between parameterizations for BDSKY and the observed data likelihood.

#### 5.2.3 Variational approximating family

We make a standard mean field assumption, which posits that members of 𝒬 completely factorize into a product of independent marginals. Letting ***ζ*** = (*R*_1_,…, *R*_*m*_, *δ*_1_,…, *δ*_*m*_, *s*_1_,…, *s*_*m*_) denote the collection of all global parameters defined above, and recalling the definition of ***z***^(*i*)^ in the preceding paragraph, we assume that

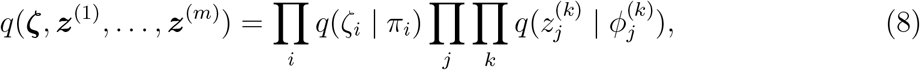

where we have introduced variational parameters *π*_*i*_ and 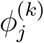 corresponding to each marginal distribution. The distributions *q*(*ζ*_*i*_ | *π*_*i*_) and 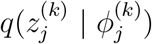 are (suitably transformed) Gaussians, so that 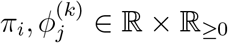 each comprises a real location parameter and non-negative scale parameter. In our model, all latent parameters, local or global, are constrained to be positive (e.g., ***R, δ***) or in the unit interval (e.g., ***s, z***^(*i*)^). For each parameter we take *q* to be an appropriately transformed normal distribution. For positive parameters we use an exponential transformation, and for parameters constrained to be in (0, 1) we use an expit (inverse logistic) transformation.

#### 5.2.4 Implementation using differentiable programming

Our Python software implementation uses automatic differentiation in order to efficiently optimize the variational objective function (Kucukelbir et al., 2017; Bradbury et al., 2018). We sample from the variational distribution and estimate the gradient of the (7) objective function with respect to the variational parameters ***π*** and ***ϕ*** using Monte Carlo integration (cf. eqn. 7). Gradients of the phylogenetic likelihood are computed in linear time using the recent algorithm of Ji et al. (2020). The complete fitting algorithm is shown in Algorithm 1.

##### Algorithm 1: Variational Bayesian Skyline (VBSKY)

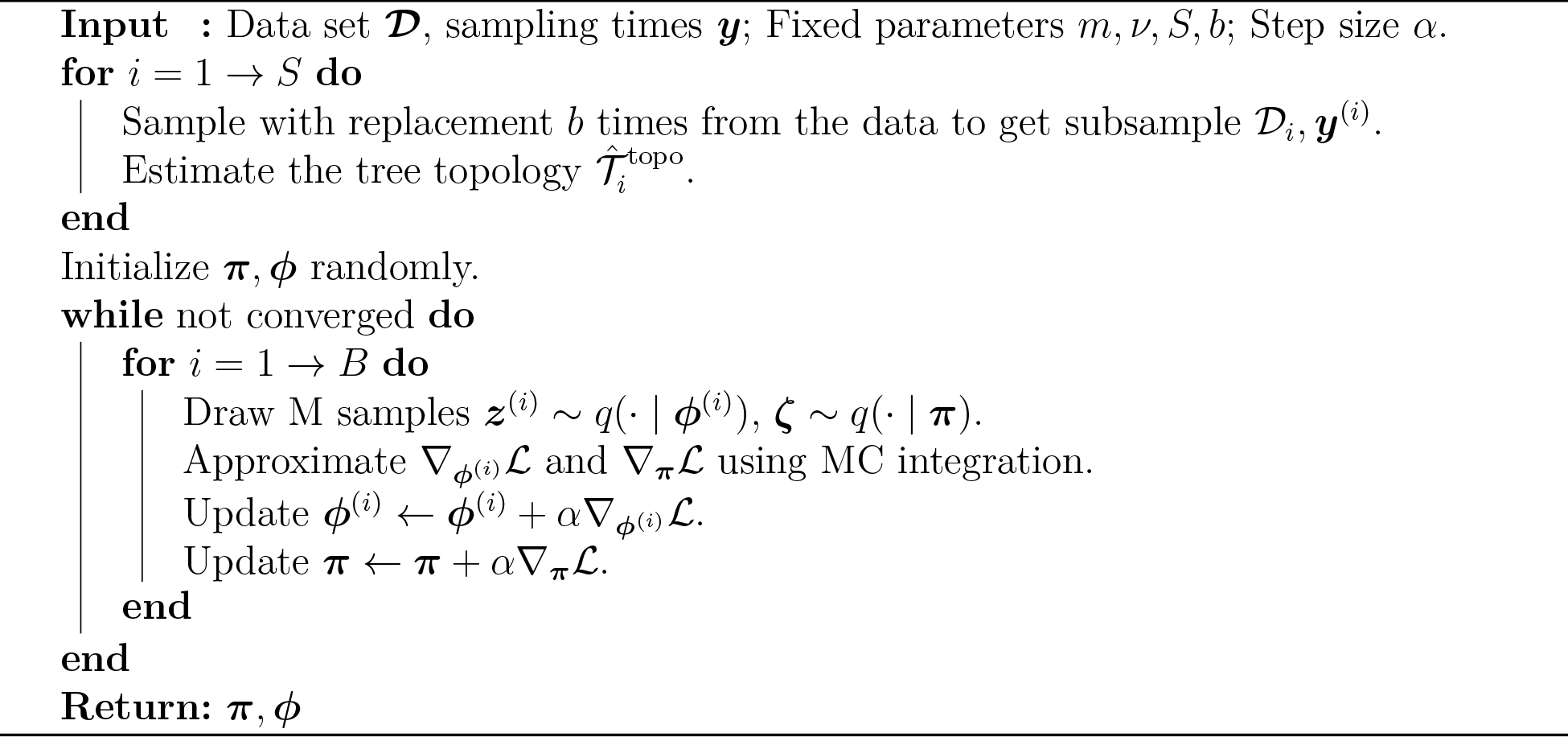

## Acknowledgments

This research was supported by the National Science Foundation (grant number DMS-2052653, and a Graduate Research Fellowship).

## Data availability

All of the data analyzed in this manuscript are publicly available. A Python implementation of our method, as well as Jupyter notebooks which reproduce our results, are located at https://github.com/jthlab/vbsky.

## Footnotes

1 At the time this manuscript was written, there were no available sequences from the Omicron variant.

## Notes

### Competing Interest Statement

The authors have declared no competing interest.

