## Supplemental materials for "Variational phylodynamic inference using pandemic-scale data"

### S-1 Birth Death Skyline

In this section we review the birth-death skyline (BDSKY) model of Stadler et al. (2013). BDSKY is a forward time model which begins with a single individual at time $t_0$ and ends at $t_m$. Throughout this section, we will refer to the start of the process as the origin. As with other skyline methods, the parameters of the model are allowed to vary over time. Specifically, given a vector $\mathbf{t} = (t_0, t_1, \ldots, t_m)$ satisfying $0 < t_1 < \cdots < t_m$, parameters are fixed between each $t_k$ and $t_{k-1}$, and allowed to vary $m$ times. The transmission rates are denoted by the vector $\boldsymbol{\lambda} \in \mathbb{R}^m_{>0}$. Similarly, the death rates are given by the vector $\boldsymbol{\mu}$ and the sampling rates by the vector $\boldsymbol{\psi}$ where each $\mu_k > 0$ and $\psi_k > 0$. In the interval $[t_{k-1}, t_k)$, every infected individual transmits at rate $\lambda_k$, recovers at rate $\mu_i$, and is sampled at rate $\psi_k$. For ease of notation, we denote $\lambda(t), \mu(t)$, and $\psi(t)$ as the transmission rate, uninfectious rate, and sampling rate at time $t$. We assume that after sampling, the individual can no longer transmit. This assumption holds in reality for many viruses as sampling is often followed by treatment or changes in behavior that would curb or limit spread. For example, those sampled with HIV would undergo antiretroviral therapy or those sampled with COVID-19 would quarantine themselves.

As described in the main text, the BDSKY model also allows for additional sampling efforts at each time $t_k$. For the reader's convenience we reproduce the notation here. All infected are sampled with rate $\rho_k$ at time $t_k$. When all sequences are sampled serially without the added sampling effort, $\rho_k = 0$ for $1 \leq k \leq m$. When all sequences are sampled contemporaneously, $\boldsymbol{\psi} = \mathbf{0}$, $\rho_k = 0$ for $1 \leq k \leq m-1$, and $\rho_m > 0$. For our work, we only consider cases where $\rho_k = 0$ for $1 \leq k \leq m-1$. We define $b_s$ as the number of sequences sampled serially, and $b_m$ to be the number of sequences sampled at time $t_m$. In other words, $b_m$ is the number of contemporaneously sampled sequences at time $t_m$. Note that $b = b_m + b_s$. The sample times of the $b_s$ serially sampled sequences are denoted by $\tilde{\boldsymbol{y}}^{(i)} = (y_1^{(i)}, \ldots, y_{b_s}^{(i)})$. Because the sequences sampled at $t_m$ have the largest sample time, $\tilde{\boldsymbol{y}}^{(i)}$ is just a truncated version of $\boldsymbol{y}^{(i)}$. When all sequences are sampled serially, $\boldsymbol{y}^{(i)} = \tilde{\boldsymbol{y}}^{(i)}$. To conserve notation, from this point onward, we will use $\boldsymbol{y}^{(i)}$ to refer to $\tilde{\boldsymbol{y}}^{(i)}$. The $b-1$ transmission event times are denoted by $\boldsymbol{x}^{(i)} = (x_1^{(i)}, \ldots, x_{b-1}^{(i)})$ where $0 < x_1^{(i)} < \cdots < x_{b-1}^{(i)}$.

The number of lineages that began before $t_k$ and are extant at $t_k$ is $n_k$. Any tree $\mathcal{T}_i$ induced by the BDSKY model is described by its tree topology $\mathcal{T}_i^{\text{topo}}$, the transmission times $\boldsymbol{x}^{(i)}$, and the sampling times $\boldsymbol{y}^{(i)}$. Letting S be the event that at we observe at least one sample, the probability density of a tree under the BDSKY model is

$$p(\mathcal{T}_i \mid \boldsymbol{\lambda}, \boldsymbol{\mu}, \boldsymbol{\psi}, \boldsymbol{\rho}, \boldsymbol{t}, S) = \frac{q_1(0)\rho_m^{b_m}}{1 - p_1(0)} \prod_{k=1}^{b-1} \lambda_{I(x_k^{(i)})} q_{I(x_k^{(i)})}(x_k^{(i)}) \prod_{k=1}^{n_s} \frac{\psi_{I(y_k^{(i)})}}{q_{I(y_k^{(i)})}(y_k^{(i)})} \prod_{k=1}^{m} q_{k+1}(t_k)^{n_k}, \quad \text{(S1)}$$

where $I(t) = k$ if $t_{k-1} \le t < t_k$, and for $k = 1, \ldots, m$ and $t_{k-1} \le t < t_k$,

$$A_k = \sqrt{(\lambda_k - \mu_k - \psi_k)^2 + 4\lambda_k\psi_k}$$
$$B_k = \frac{(1 - 2(1 - \rho_k)p_{k+1}(t_i))\lambda_k + \mu_k + \psi_k}{A_k}$$
$$p_k(t) = \frac{\lambda_k + \mu_k + \psi_k - A_k \frac{e^{A_k(t_k - t)}(1+B_k)-(1-B_k)}{e^{A_k(t_k - t)}(1+B_k)+(1-B_k)}}{2\lambda_k}$$
$$q_k(t) = \frac{4e^{-A_k(t-t_k)}}{(e^{-A_k(t-t_k)}(1 + B_k) + (1 - B_k))^2},$$

and $p_{m+1}(t_m) = 1$.

### S-2 Estimating the tree topology

In this section we explain how we estimate the tree topology $\hat{\mathcal{T}}_i^{\text{topo}}$ for each subsample $\mathcal{D}_i$. We employed a simple heuristic method by fitting serial-sample unweighted pair grouping method with arithmetic means (sUPGMA) (Drummond et al., 2000). As the name alludes to, sUPGMA is a tree reconstruction algorithm based on the unweighted paired group method with arithmetic means (UPGMA) (Sneath and Sokal, 1983).

We first describe the UPGMA algorithm followed by the sUPGMA algorithm. Both algorithms require a pairwise distance matrix. Taking the sequences of our subsample, $\mathcal{D}_i \in \{A, C, G, T\}^{b \times L}$, we simply take the Hamming distance between all $\binom{b}{2}$ pairs of sequences. That is for a given pair of sequences $s, t \in \mathcal{D}_i$, the distance is $d(s,t) = \sum_{k=1}^{L} \mathbb{1}_{(s_k \neq t_k)}$. For clusters, the mean distance between each element in the clusters. That is

$$d(A, B) = \frac{1}{|A||B|} \sum_{s \in A} \sum_{t \in B} d(s, t),$$

where $A$ and $B$ both represent clusters. At each step the two clusters with the smallest distance between them are combined into a new cluster, and the distances are recalculated between the newly formed cluster and the other clusters. Clusters can be made up of a single sequence. We start with $b$ clusters initially and reduce the number of clusters by one at each step. This is repeated until there is only one single cluster.

While we could naively use UPGMA to get our tree topologies, because UPGMA does not account for sample times, it is possible the algorithm would give us topologies that are impossible given the sample times. We use sUPGMA to ensure that the estimated topology is not only possible, but realistic. Consider our subsample of sample times $\boldsymbol{y}^{(i)} = \{y_1^{(i)}, y_2^{(i)}, \ldots, y_b^{(i)}\}$. Let $d(u_i, v_j)$ be the distance between $i$th sequence with sample time $y_u^{(i)}$ and the $j$th sequence with sample time $y_v$. We assume that $u < v$. and model $d(u_i, v_j)$ by its expectation, $\mathbb{E}(d(u_i, v_j)) = \Theta_u + \omega(y_v^{(i)} - y_u^{(i)})$, where $\Theta_u$ is the expected average distance between any two sequences at time $y_u^{(i)}$, and $\omega$ is the expected number of substitutions per unit time. The procedure for sUPGMA is as follows.

1. Estimate the set of parameters $\{\Theta_1, \ldots, \Theta_q, \omega\}$ using regression:

$$d(u_i, v_j) = \sum_{k=1}^{q} \Theta_k X_k + \omega(y_v^{(i)} - y_u^{(i)}) + \epsilon,$$

   where $X_k = 1$ if $k = u$ and $k = 0$ otherwise.

2. Correct the original pairwise distances

$$c(u_i, v_j) = d(u_i, v_j) + \omega(y_u^{(i)} + y_v^{(i)} - 2y_1^{(i)}).$$

3. Cluster using the UPGMA algorithm as described earlier.

We note that while the complete sUPGMA algorithm returns both the tree topology and the branch lengths, we only use this procedure to obtain the tree topology. As branch lengths are continuous variables, we will estimate those using stochastic variational inference. Although there are maximum likelihood based tree reconstruction methods we could use such as IQ-TREE (Minh et al., 2020), since we are only concerned with the topology rather than the entire tree, we prioritize the faster algorithm.

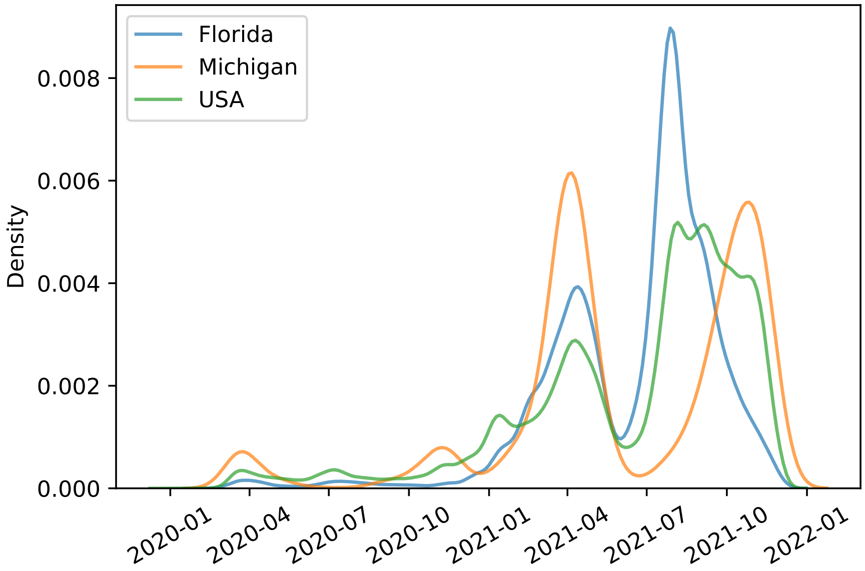


**Figure S1:** Distribution of sample times for Florida, Michigan, and the USA.

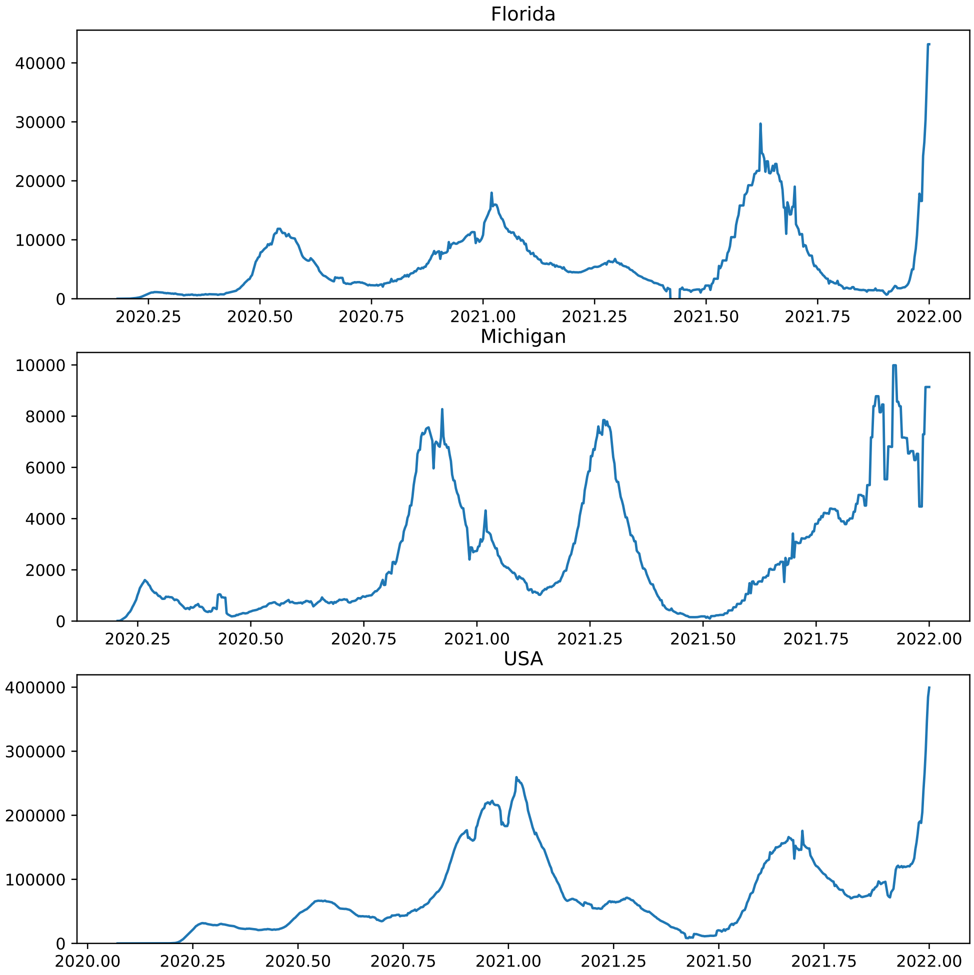


**Figure S2:** Daily new cases of COVID-19 over time for Florida, Michigan, and the USA.

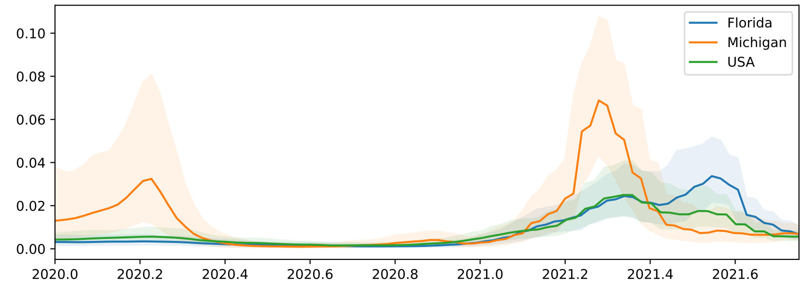


**Figure S3:** The posterior median and equal-tailed 95% credible interval of $s$ for Florida, Michigan, and the USA using an uninformative smoothing prior.

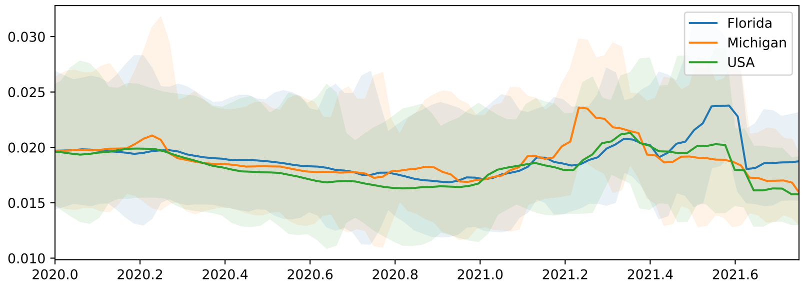


**Figure S4:** The posterior median and equal-tailed 95% credible interval of $s$ for Florida, Michigan, and the USA using less smoothing.

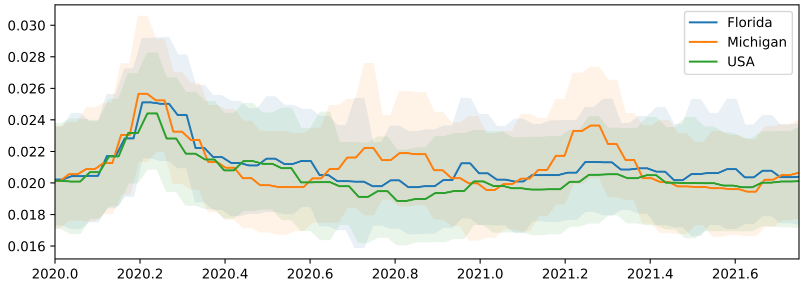


**Figure S5:** The posterior median and equal-tailed 95% credible interval of $s$ for Florida, Michigan, and the USA using biased sampling.

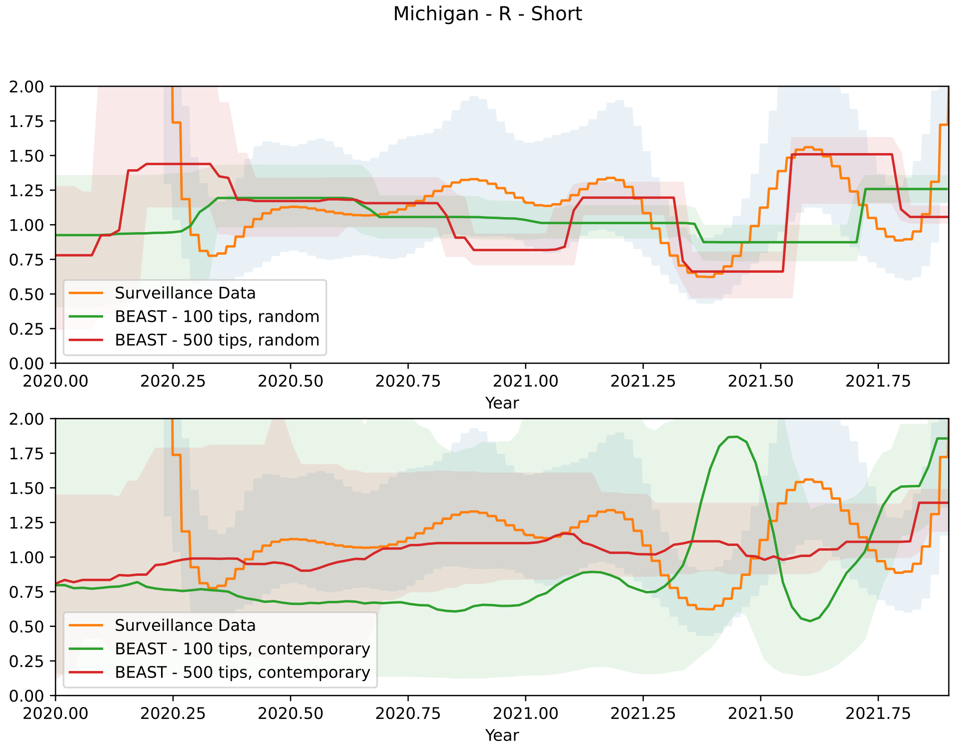


**Figure S6:** The posterior median and equal-tailed 95% credible interval of $R$ for Michigan given by BEAST. The sampler was allowed to run as long as it VBSKY to analyze the Michigan data. This is referred to as the short run in the text.

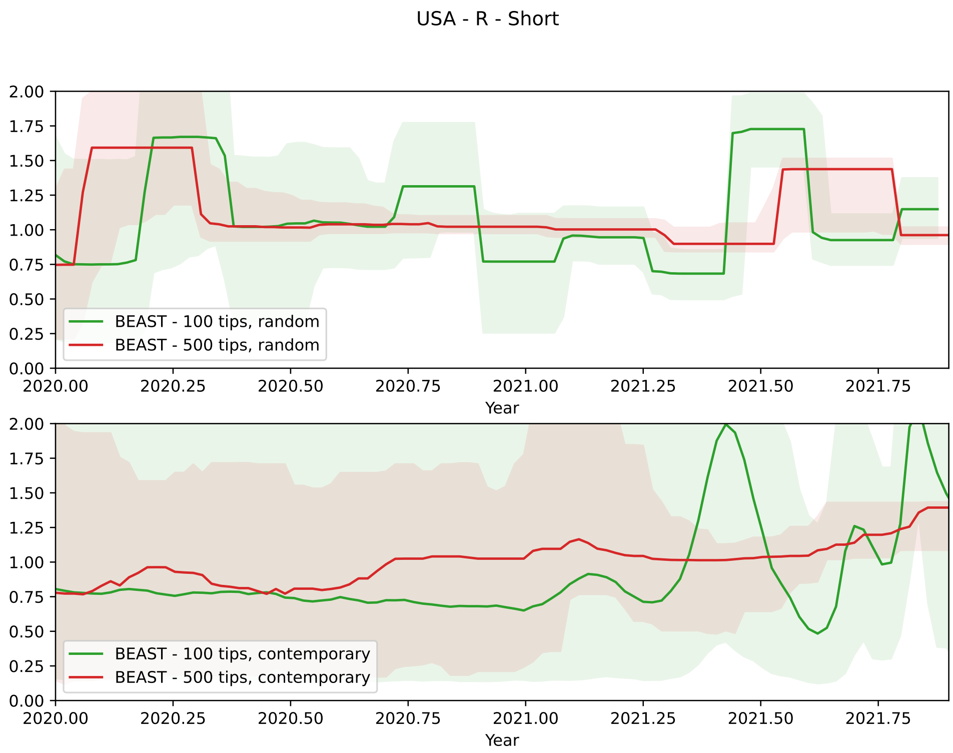


**Figure S7:** The posterior median and equal-tailed 95% credible interval of $R$ for the USA given by BEAST. The sampler was allowed to run as long as VBSKY to analyze the USA data. This is referred to as the short run in the text.

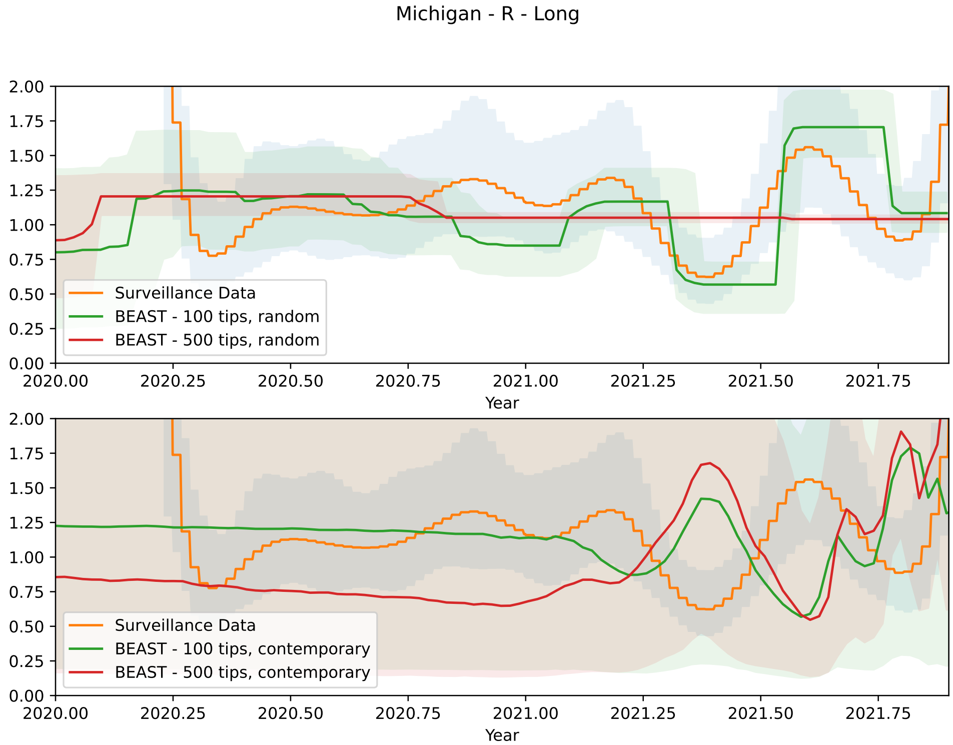


**Figure S8:** The posterior median and equal-tailed 95% credible interval of $R$ for Michigan given by BEAST. The sampler was allowed to run for 100 million steps or 24 hours. This is referred to as the long run in the text.

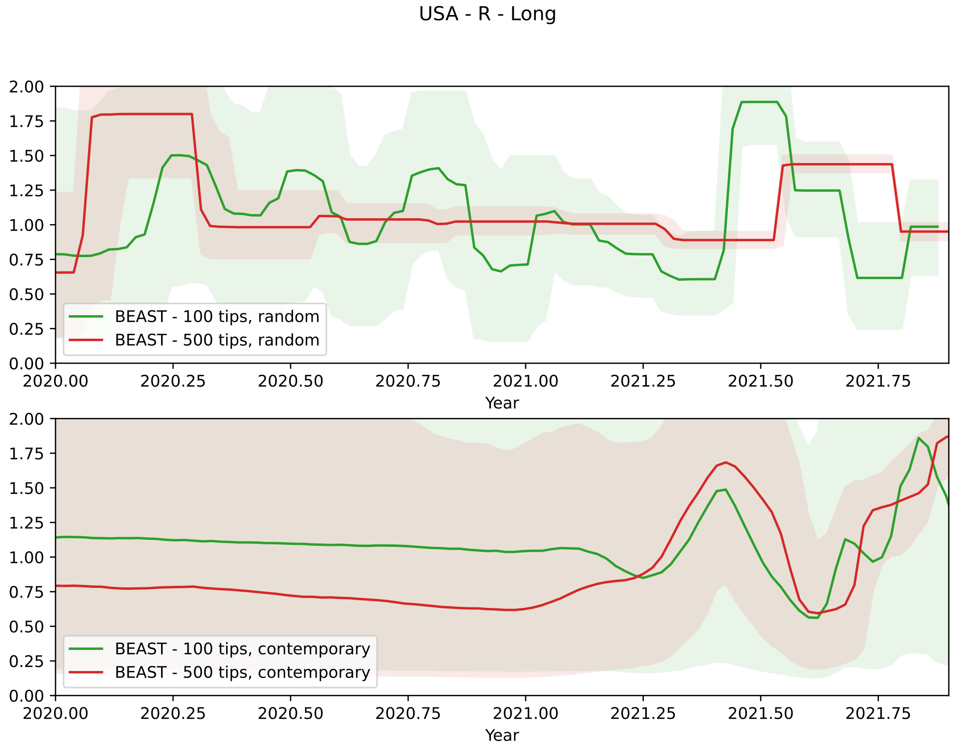


**Figure S9:** The posterior median and equal-tailed 95% credible interval of $R$ for the U.S. given by BEAST. The sampler was allowed to run for 100 million steps or 24 hours. This is referred to as the long run in the text.

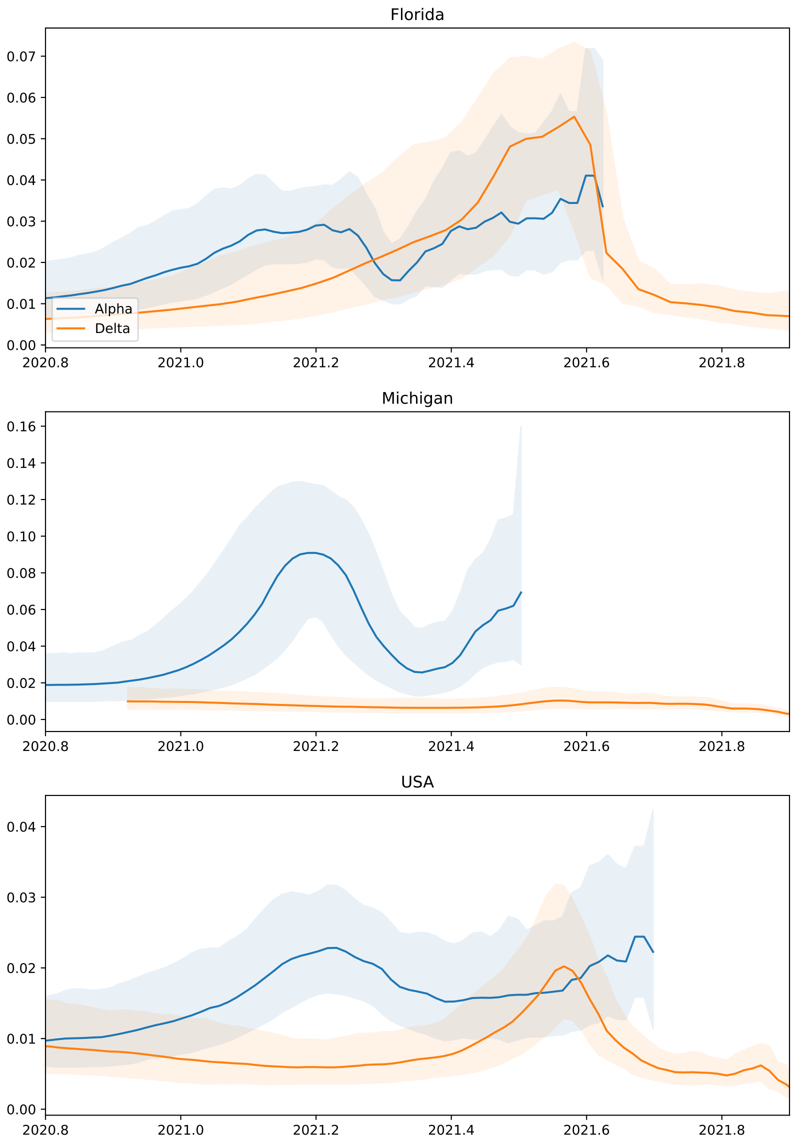


**Figure S10:** The posterior median and equal-tailed 95% credible interval of $s$ for the Alpha and Delta variants.

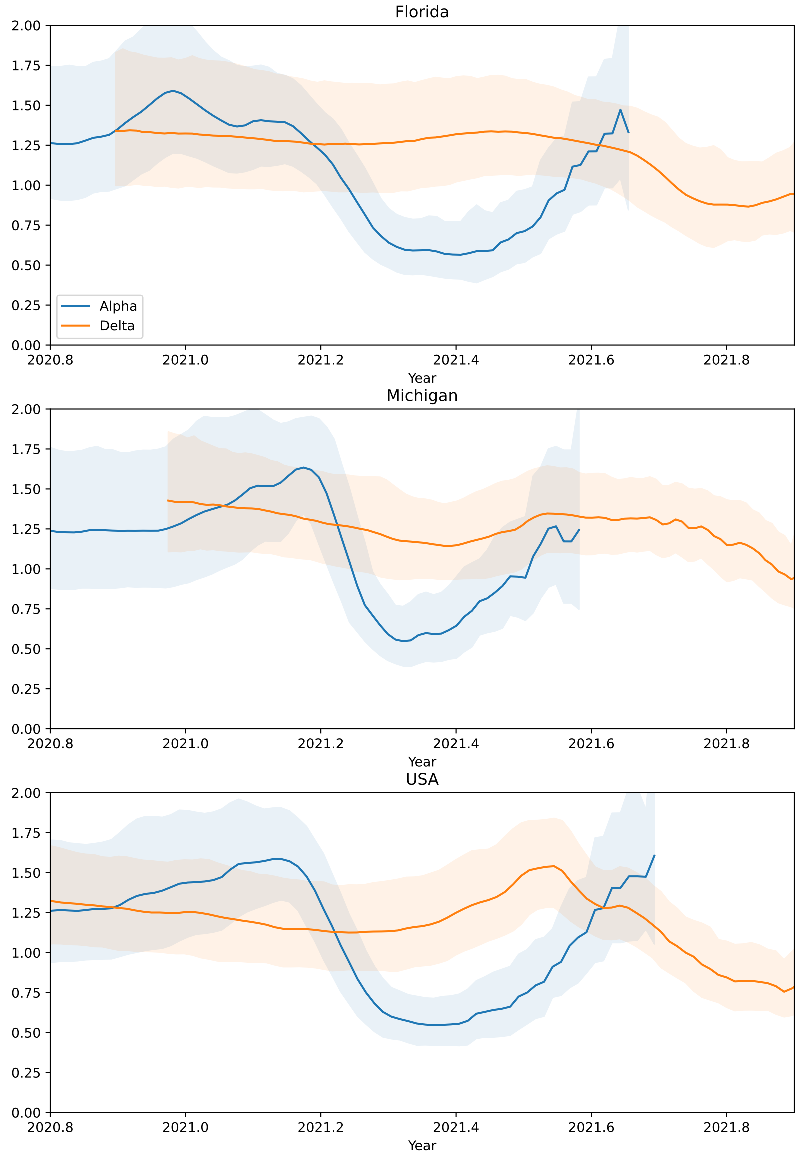


**Figure S11:** The posterior median and equal-tailed 95% credible interval of $R$ for the Alpha and Delta variants.

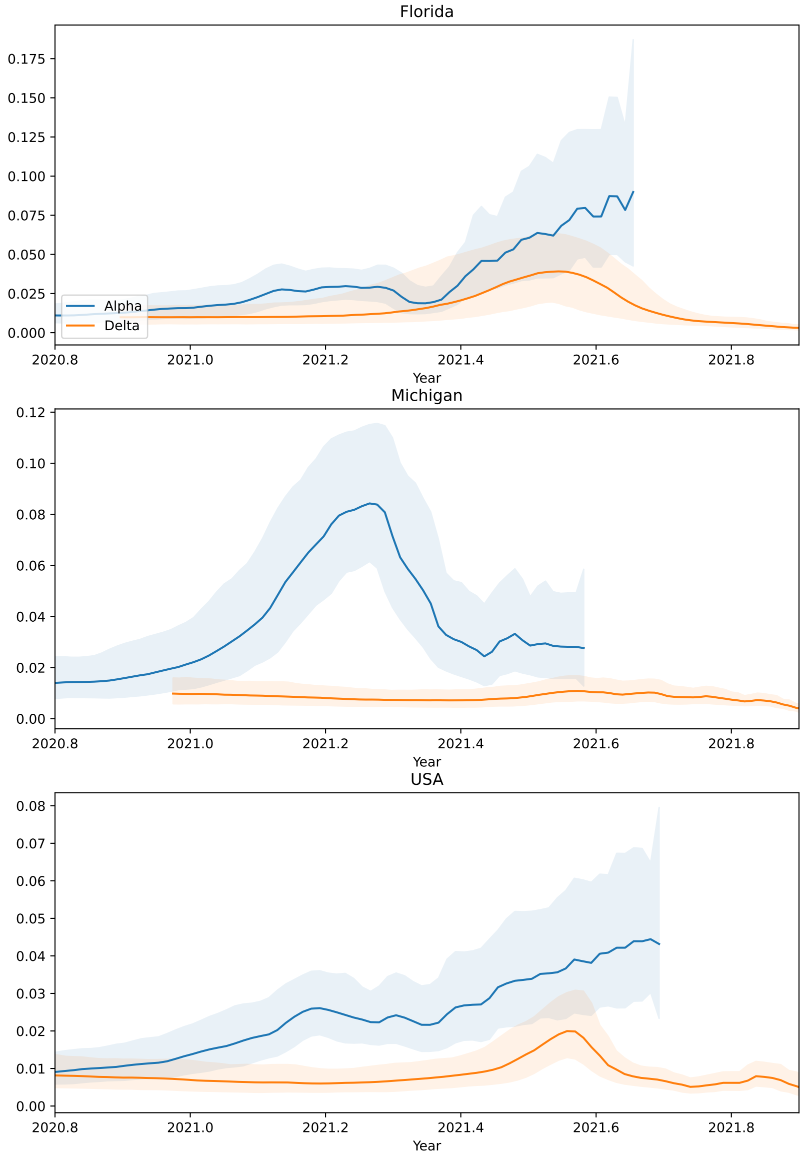


**Figure S12:** The posterior median and equal-tailed 95% credible interval of $s$ for the Alpha and Delta variants.
